# A unified *in vitro* to *in vivo* fluorescence lifetime screening platform yields amyloid β aggregation inhibitors

**DOI:** 10.1101/2022.03.28.485913

**Authors:** Súil Collins, Liisa van Vliet, Fabrice Gielen, Matej Janeček, Sara Wagner Valladolid, Chetan Poudel, Giuliana Fusco, Alfonso De Simone, Claire Michel, Clemens F. Kaminski, David R Spring, Florian Hollfelder, Gabriele S Kaminski Schierle

**Author notes:** **Correspondence and requests for materials** should be addressed to Florian Hollfelder, Gabriele S Kaminski Schierle or David Spring.

## Abstract

Inhibiting the aggregation of amyloid β (1-42) is a promising strategy for the development of disease-modifying Alzheimer’s disease therapeutics. To date, however, no sufficiently efficacious inhibitors have been identified, despite the best efforts of >200 advanced drug development campaigns. This failure can be attributed to limitations in current compound screening and *in vivo* validation assays. Here, we report an *in vitro* to *in vivo* screening platform based on the use of a fluorescence lifetime aggregation sensor. The microfluidic “nanoFLIM” assay developed circumvents issues that plague conventional assays, such as lack of reproducibility, high cost and artefactual false read-outs. The fluorescence lifetime sensor can also dynamically monitor peptide aggregation in cellular and *Caenorhabditis elegans* disease models, providing directly comparable aggregation kinetics, which is not achievable by any other method. The power of this unified system for accelerating hit-to-lead strategies, lowering attrition rates and expediting *in vivo* screening, was demonstrated with a pilot screening campaign of 445 compounds, revealing a new inhibitor that can inhibit amyloid β self-assembly *in vitro* as well as in cellular and whole organism disease models.

## Introduction

Aggregation of the peptide amyloid β (1-42) (Aβ_42_) is a pathological hallmark of Alzheimer’s disease (AD), a multifaceted neurodegenerative disorder for which no preventative measures or disease modifying therapies exist.^1, 2^ The development of Aβ_42_ aggregation inhibitors has long been recognised as a potential strategy for AD treatment,^3, 4, 5, 6^ but to date only one inhibitory drug has been approved, subject to an unusual nine-year post-approval trial.^7, 8^ Indeed, more than 200 unsuccessful attempts to develop medicines in clinical trials to treat and potentially prevent AD are on record,^9, 10^ suggesting that finding therapeutic molecules in this area is much harder than in other drug development campaigns. However, the number of drugs candidates in clinical trials at any time remains above 100.^11^ Setbacks in the development of such clinical candidates stem from a variety of issues along the drug development pathway, including the types of assays historically used and the difficulty of integrating biophysical and cell-based studies.^12^ The intrinsically disordered nature of the peptide and aggregate heterogeneity has limited structural studies on the peptide, thereby occluding structure-based design strategies.^13, 14, 15^ Available compound libraries often lack novelty and scaffold diversity,^16^ and current screening approaches are limited by reagent consumption, assay reproducibility and spectral inference from intrinsic properties of the test compounds. A major limitation in the development of AD therapeutics is the absence of methods to test the activity of hit compounds in cellular or whole organism models quickly and reliably: there are currently no high-throughput screening systems to directly monitor cellular Aβ_42_ aggregation in real time, or compare in vitro and cellular anti-aggregation activity.^17^ Current techniques rely on fixing cells to stain amyloid deposits or simple cell viability tests, neither of which are dynamic or revealing about the underlying aggregation processes. Filtering and prioritising *in vitro* hits for those that show the strongest *in vivo* activity is often too laborious to pursue extensive ‘hit-to-lead’ strategies, potentially resulting in the advancement of lead compounds whose sub-optimal *in vivo* activity only becomes apparent in late stage development.^18, 19, 20^

Herein, we report an approach that addresses these issues by employing a unified assay format to cover *in vitro* to *in vivo* compound screening for Aβ_42_ aggregation inhibitory activity, in order to better identify and validate drug candidates, and to avoid wasting efforts on false positive results that have previously led to failed drug discovery campaigns. This comprehensive assay is based on the use of an amyloid aggregation fluorescence lifetime sensor, whereby Aβ_42_ aggregation is monitored by changes in the fluorescence lifetime of an attached fluorophore, which is significantly quenched upon self-assembly of the peptide.^21, 22^ Changes in the aggregation profile in the presence of a small molecule is indicative of a modulatory effect on Aβ_42_ self-assembly, be it to promote, delay or inhibit the process.

A medium-throughput microfluidic assay (dubbed the *nanoFLIM*) was designed to screen compound libraries using fluorescence lifetime imaging microscopy (*FLIM*), with large sample sizes (>100 assays per experiment) and *nanolitre* volume requirements (18 nL per test). To demonstrate the potential of the system, a selection of novel chemical libraries – developed using diversity-oriented synthesis (DOS) or rationally targeted drug discovery strategies – was interrogated, yielding **MJ040**, a novel lead Aβ_42_ aggregation inhibitor. The fluorescence lifetime sensor protocol was also applied to dynamically screen compounds for Aβ_42_ anti-aggregation activity in SH-SY5Y neuroblastoma cells, where **MJ040** was again shown to display an inhibitory effect. The activity of a rationally designed prodrug version, **MJ040X**, was further validated by probing *in vivo* Aβ_42_ aggregation in the disease model *Caenorhabditis elegans*, in which treatment was seen to delay or completely inhibit the aggregation process.

In addition to an unrivalled economy in reagent consumption, the *nanoFLIM* platform provides the only unified direct comparison of Aβ_42_ aggregation propensity in the presence of small molecules *in vitro*, in live cells and in a whole organism disease model. The successful identification of **MJ040X** as a novel lead Aβ_42_ aggregation inhibitor, capable of preventing amyloid formation in all three relevant formats, suggests that promising candidates for the development of therapeutically active AD treatment readily result from this approach, thereby providing a route to drug development for a hitherto elusive challenge.

## Results and discussion

### Small molecule library screening at the nanoliter scale identifies novel inhibitory scaffold

To overcome problems that have plagued previous screening campaigns, namely assay variability, data quality and reagent consumption,^12, 23^ a novel workflow was designed in which Aβ_42_ aggregation was monitored within arrayed nanoliter droplets using fluorescence lifetime imaging.^24, 25^ The arrayed format enabled the simultaneous imaging of 110 droplets, each corresponding to a distinct experiment. **Figure 1a** shows an overview of the nanoFLIM assay platform, in which 110 traps sit along a serpentine channel in a microfluidic chip. The droplets (18 nL) lodged in these traps were generated and deposited using droplet-on-demand technology,^26, 27^ producing defined quantities of partially labelled monomeric peptide (845 pg per droplet) and the respective drug candidate molecule (360 pg per droplet) in a precisely known order (**Fig. 1a,b**). Once droplets were sequentially trapped in the device, reaction progress was imaged over several hours, to give characteristic aggregation profiles (**Fig. 1c**). 445 compounds, with 5-10 repeats per compound at a single concentration, were screened to give start point and end point values, taking only ~70 hours for ~2400 experiments. The compounds originated from novel chemical libraries synthesised by diversity-oriented approaches or specifically designed to incorporate medicinally relevant features (**Supplementary Results, Supplementary Table 1**). A sample screen showing the fluorescence lifetime change of trapped Aβ_42_ droplets with 9 repeats of 9 compounds is shown in **Figure 2a**. Such screening efforts resulted in the identification of hit compound **MJ040** (**Fig 2b**), which originates from a cinchophen scaffold library and exerts a profound inhibitory effect on Aβ_42_ aggregation compared to other library members (**Fig. 2a**).^28^ The ability of **MJ040** to inhibit Aβ_42_ aggregation was quantified by nanoFLIM measurement of Aβ_42_ aggregation kinetics at several concentrations of the compound, giving an IC_50_ value of 3.8 ± 0.8 μM (**Fig. 2c,d**). Notably, we multiplexed aggregation assays to overcome batch to batch variability even shown for recombinant Aβ^29^ by carrying out repeats. The high number of repeats in this assay made it possible to obtain reliable aggregation data immediately from one set of measurements, with incomparably low reagent consumption. The low micromolar range of the IC_50_ value is similar to known *in vitro* inhibitory compounds, e.g. EGCG (IC_50_ 6.4 ± 0.7 μM),^30^ an inhibitor that has reached phase III clinical trials (NCT00951834) for its Aβ_42_ anti-aggregation activity. Two compounds, **MJ001** and **MJ042** (**Fig. 2a,b**), showed weaker, yet significant inhibitory activity and were carried forward together with **MJ040** for further validation.

**Figure 1.**
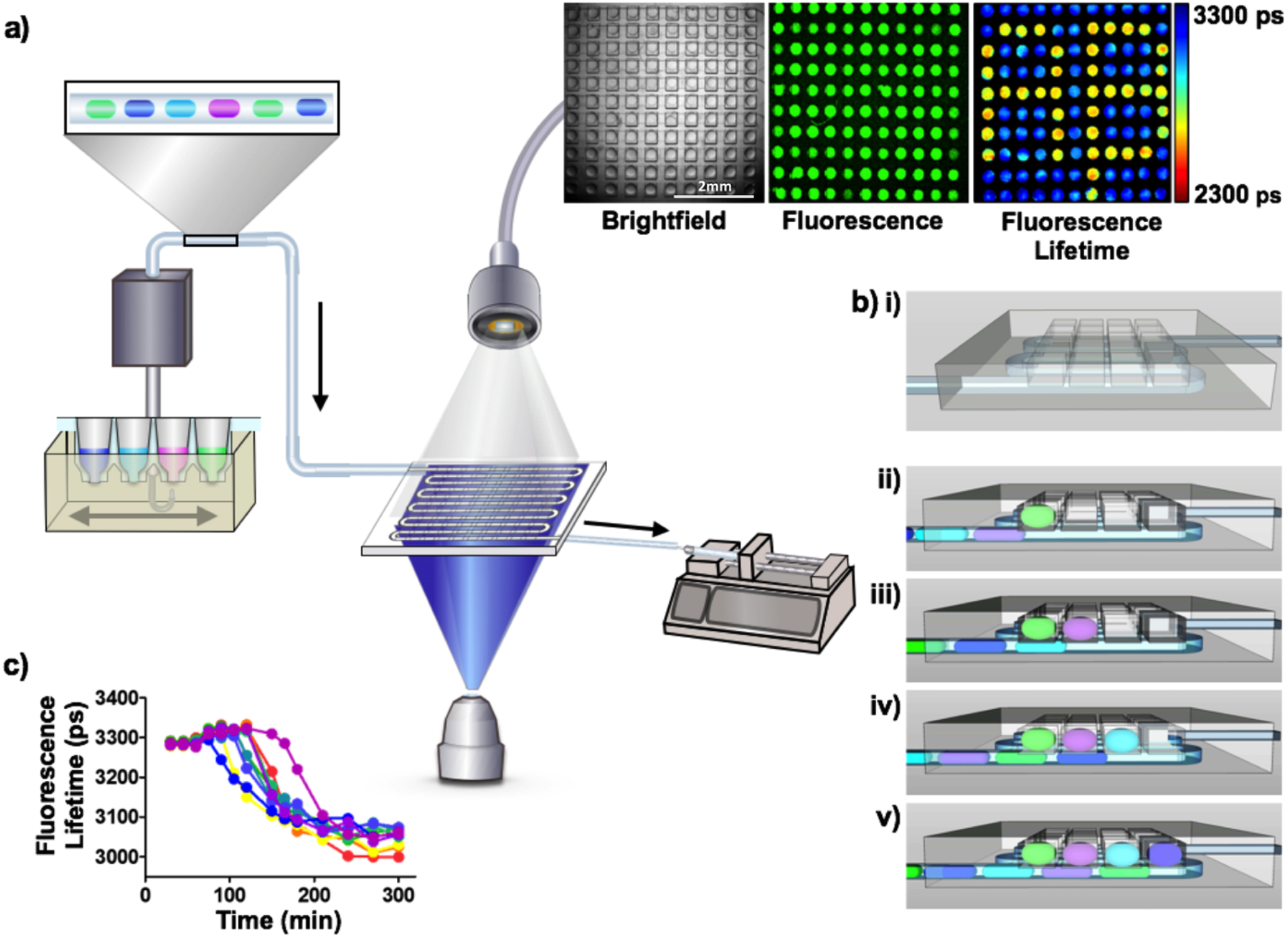
Overview of the nanoFLIM. a) Assay workflow. Using droplet-on-demand technology (Mitos Dropix, Dolomite^26^) droplets containing monomeric peptide together with test compounds are formed and filled into a droplet array chip in a predefined order. The brightfield image insert shows the filled microfluidic device, in which 110 droplets, with 18 nL of aqueous solution per droplet, are trapped within square grids above a serpentine channel (Scale bar = 2 mm). Fluorescence lifetime imaging is used to monitor the aggregation kinetics of the peptide, which is partially labelled with a reporter dye, and the effect that different extrinsic factors or inhibitory compounds can have on the process. The fluorescence intensity of monomeric or aggregated peptide droplets are similar. However, the fluorescence lifetime sensor provides a quantitative measure of the aggregation state of the peptide, which are either monomeric (blue, lifetime ~3200 ps) or aggregated (yellow, lifetime ~2700 ps). The ability of the system to monitor the process of Aβ_42_ aggregation with different peptide labelling densities, under various pH, temperature, shearing conditions and with a variety of known inhibitory small molecules is shown in **Extended Data Figure 3, and Supplementary Figures 1-4. b)** *Schematic of microfluidic chip filling*. ***i)*** Empty device showing square grids aligned over a serpentine channel. ***ii)*** Droplets of a known sequence fill into the serpentine channel and travel until they reach an empty grid, where ***iii)*** the leading droplet floats upwards and is trapped in the first grid and held for several hours for imaging. ***iv)*** Subsequent droplets flow under previously trapped droplets until ***v)*** they reach an empty grid where they become trapped. **c)** Aβ_42_ aggregation profiles for 10 droplets formed from the same 10 μL stock solution of peptide. A great discrepancy between the initiation of aggregation in the first droplet and the last is observed, highlighting the need for many repeats when monitoring self-assembly of this highly aggregation prone peptide. 20 μM Aβ_42_-488, 20% labelled.

**Figure 2:**
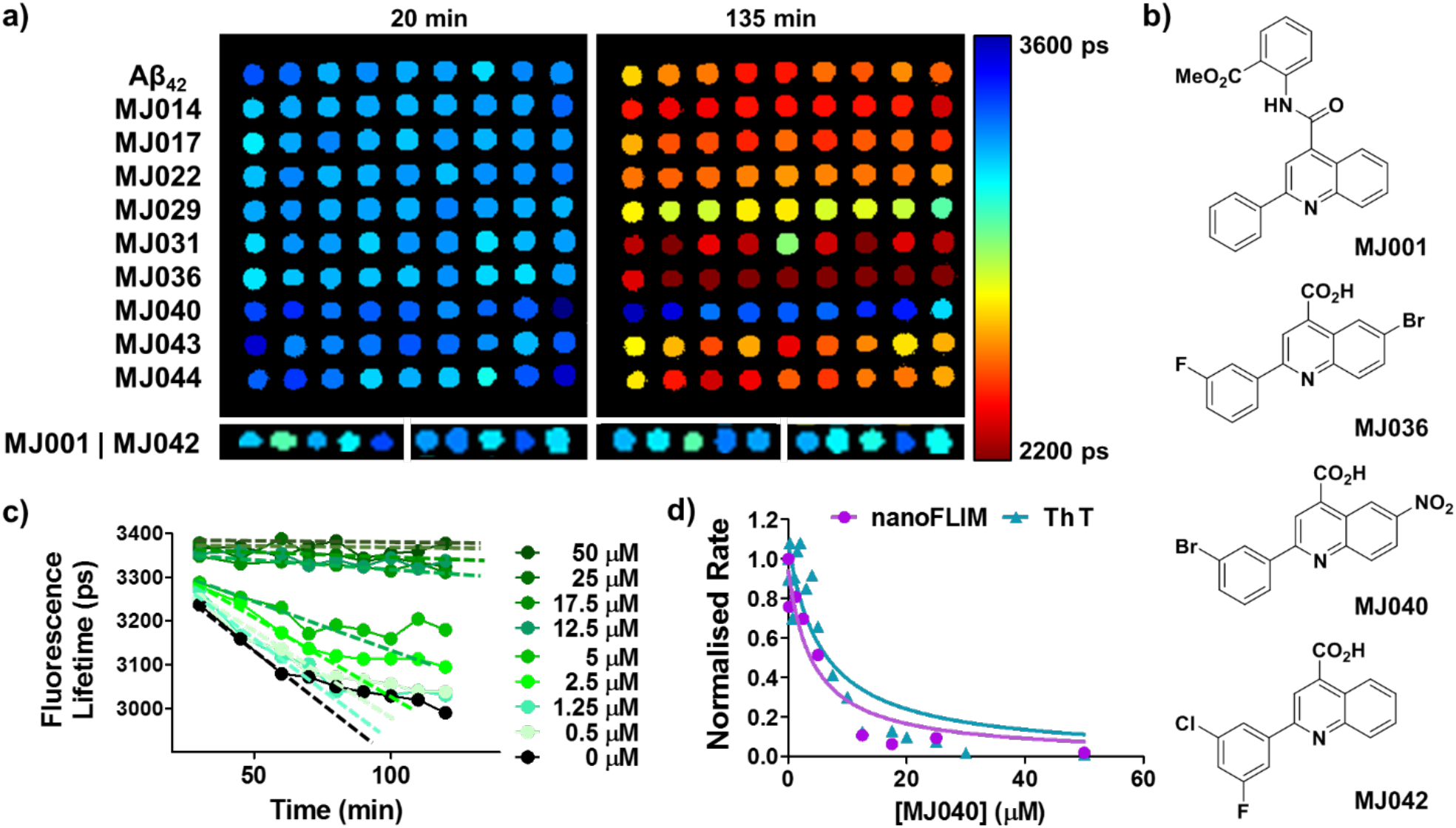
Screening of compound collections using the nanoFLIM identifies active inhibitory compounds. **a)** Example of the nanoFLIM screening approach for hit identification. The raw FLIM images show rows of nine droplet replicates containing peptide and test compounds at start (20 min., left) and end (135 min, right) time points. The changes in fluorescence lifetime are monitored to give averaged aggregation profiles, as shown in panel c). Compounds capable of preventing the lifetime change associated with peptide aggregation, e.g. **MJ040** (row 8) and **MJ001** and **MJ042** (inserts) are selected for further study. 10 μM Aβ_42_-488, 50% labelled, 10 μM small molecule, droplet diameter = 200 μm. (Details of the FLIM analysis and representative photon counts are shown in **Extended Data Figure 4**. The full cinchophen library and associated nanoFLIM data are shown in **Supplementary Figure S5 & S6**.) **b)** Structures of the hit compounds from the cinchophen library carried forward for further characterisation. **c)** Aβ_42_ aggregation profiles obtained by filling the device with a concentration gradient of **MJ040** (n = 9). **d)** IC_50_ graphs using the initial rates obtained for a concentration gradient from the nanoFLIM and ThT fluorescence assays.^31^ nanoFLIM IC_50_ = 4.3 ± 1.3 μM, ThT fluorescence IC_50_ = 5.8 ± 1.7 μM. The lines are fits to the equation: 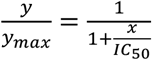

### Inhibitory activity of hit compounds validated by conventional Thioflavin T assays and structural analysis

Thus far, screening for Aβ_42_ aggregation inhibitors has mainly employed Thioflavin T (ThT) fluorescence assays, whereby a red shift and enhancement in the fluorescent signal of the ThT dye upon binding to structures rich in β-sheets is correlated with increasing protein aggregation.^32^ The anti-aggregation activity observed using the ThT fluorescence assay and the nanoFLIM was compared, and the data for four notable compounds are shown in **Figure 3**. Both the nanoFLIM (**Fig. 3a**) and ThT (**Fig. 3b**) assays indicated that **MJ040** exerts a strong inhibitory effect, with calculated IC_50_ values of 4.3 ± 1.3 μM and 5.8 ± 1.7 μM, respectively (**Fig 2d**). The mode of interaction of MJ040 was investigated by a combination of NMR spectroscopy and molecular modelling, allowing us to locate the MJ040 binding site in a hydrophobic cleft near the C-terminus of monomeric Aβ_42_ (**Extended Data Figure 1**).

**Figure 3:**
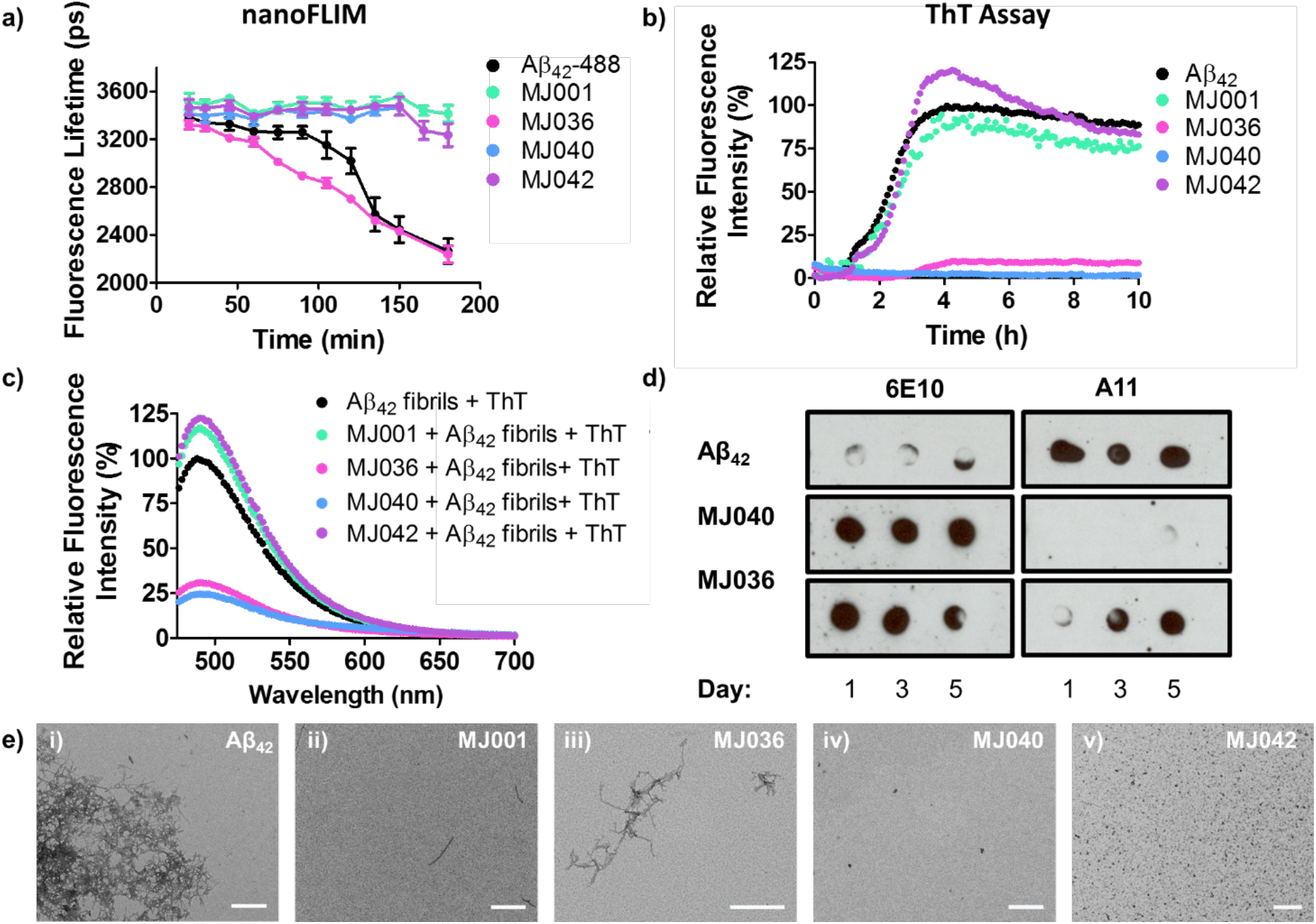
Comparison of the inhibitory activity detected using the nanoFLIM, ThT fluorescence assays, A11 dot blots and TEM imaging. **a)** NanoFLIM Aβ_42_ aggregation profiles in the presence of select compounds. **MJ001, MJ040** and **MJ042** exert an inhibitory effect, but aggregation is seen to proceed in the presence of **MJ036**. 10 μM Aβ_42_-488, 50% labelled, 50 μM compound. **b)** Aβ_42_ aggregation curves with the addition of the select compounds, as monitored by ThT fluorescence. The curve for Aβ_42_ (black) represents the time course of Aβ_42_ aggregation in the absence of inhibitors, the plateau of which is taken to represent fibril mass concentration and is set as 100%.^7, 30^Compounds **MJ040** and **MJ036** show a strong inhibitory effect. **MJ001** does not appear to perturb the aggregation process and the profile of **MJ042** is overly fluorescent relative to that of Aβ_42_ alone. The relative ThT fluorescence data for the full cinchophen library is shown in **Supplementary Figure 9**. 10 μM Aβ_42_, 20 μM ThT, 50 μM compound, ex = 440 nm, em = 488 nm. **c)** Emission spectrum of pre-formed Aβ_42_ fibrils and ThT samples in the presence of select compounds (ex = 440 nm), expressed relative to that of fibrillary Aβ_42_ and ThT alone (peak set 100%). **MJ040** and **MJ036** significantly quench the ThT fluorescence in the presence of fibrils. **MJ001** and **MJ042** are intrinsically fluorescent at the emission wavelength used to monitor aggregation in the ThT fluorescence assay (488 nm), which may mask the inhibitory activity of the compounds in the ThT fluorescence aggregation screen. The relative emission data for the full cinchophen library in the presence of preformed fibrils at 440 nm excitation is shown in **Supplementary Figure 10**, and **Supplementary Figure 11** shows the absorbance spectra for the compounds relative to Aβ_42_ fibrils with **d)** Dot blot time course investigating the formation of potentially toxic species, using oligomer-specific antibody A11.^33^ The 6E10 control antibody, which detects all Aβ_42_ species, shows a positive result for all samples. Oligomeric structures were present following 1, 3 and 5 days of incubation in the absence of any compounds or in the presence of **MJ036**. No A11 sensitive species were detected following incubation with **MJ040. e)** TEM images of the Aβ_42_ species formed following 7-day incubation with the compounds of interest. ***i)*** Aβ_42_ control; ***ii)* MJ001**; ***iii)* MJ036**; ***iv)* MJ040**; ***v)* MJ042**. In all cases nanoFLIM screening correctly predicted if the compound had a modulatory effect on the aggregation process. The ThT assay incorrectly identified one false positive (**MJ036**) and two false negatives (**MJ001** and **MJ042**). Scale bar = 500 nm.

With compound **MJ036**, conflicting results were obtained in the two assay formats. In the fluorescence lifetime sensor measurements, no inhibitory activity was observed with **MJ036** (**Fig. 3a**), while the ThT fluorescence assay identified **MJ036** as a potent aggregation inhibitor (**Fig. 3b**). To resolve this apparent inconsistency, morphological analysis of the aggregation products by transmission electron microscopy (TEM) (**Fig. 3e**) and atomic force microscopy (AFM) (**Supplementary Fig 7**) was performed. These orthogonal techniques rule out non-spectral and fluorescence interference, providing a method of assessing fibril formation independent of the presence of extrinsic dyes. Many aggregated species were observed following 7-day incubation with **MJ036** (**Fig. 3c-iii**), and only very few with **MJ040** (**Fig. 3c-iv**). This suggests that **MJ036** had been incorrectly assigned as a strong inhibitor by ThT fluorescence. Based on nanoFLIM it can be correctly discounted as a *false positive*. Dot blot assays, with the toxic oligomer-specific A11 antibody, were also used to investigate the formation of aggregated species Aβ_42_ in the presence of the compounds (**Fig. 3d**).^33^ A11-sensitive species were not detected in the **MJ040** treated sample, suggesting that the small structures observed by the scanning microscopy techniques were innocuous aggregates. In contrast, A11-immunoreactive structures were detected in the **MJ036** treated sample. Thus, **MJ036** neither prevents the formation of Aβ_42_ fibrils nor potentially toxic oligomeric Aβ_42_ species, and was therefore incorrectly assigned as an inhibitor by ThT fluorescence analysis.

To further assess the inadequacy of the conventional ThT assay^34^ for screening the cinchophen library, the fluorescence emission spectra of ThT with previously formed Aβ_42_ fibrils in the presence of the compounds was investigated (**Fig. 3c)**. The addition of either **MJ036** or **MJ040** was shown to reduce the ThT emission signal (with 440 nm excitation, as typically used in the real time assay), indicating that the compounds interfere with the ThT assay readout as a result of their intrinsic fluorescence properties or competitive binding interactions with the peptide or ThT dye itself.^35^ As such, the inhibitory activity of these compounds cannot be reliably assessed by means of a ThT fluorescence assay. A reduction in fluorescence intensity was not observed upon addition of **MJ036** and **MJ040** at 480 nm excitation, the wavelength used for fluorescence lifetime imaging (**Supplementary Figure 8)**. The intrinsic spectroscopic properties of **MJ001** and **MJ042** are also believed to contribute to incorrect assignments in the ThT fluorescence assay, in this case *false negative* results. When added to the ThT-Aβ_42_ fibril sample, the emission at 488 nm (the emission wavelength used in the conventional ThT assay) is higher than that of the Aβ_42_ fibrils and dye alone, thereby potentially masking the compounds inhibitory activity against Aβ_42_ aggregation **(Fig. 3b,d**), which may explain why they will *not* be found in a screen based on ThT fluorescence. The inhibitory activity of **MJ001** and **MJ042** was validated using TEM, where in both cases only small aggregates were observed following compound treatment (**Fig. 3c-ii,-iv**). NanoFLIM screening, therefore, is less susceptible to misleading readouts that produce false positive and negative results than the conventional ThT assay when screening spectroscopically active small molecule libraries.

### Prodrug synthesis generates a cell active inhibitory lead compound

To investigate the ability of lead **MJ040** to inhibit Aβ_42_-induced cell death, MTT cell vitality tests were performed. SH-SY5Y cells were employed, as this human-derived neuroblastoma cell line displays many biochemical and functional features of human neurons and has consequently seen widespread use as a human neuronal cell model in cytotoxicity assays.^33, 36, 37^ Monomeric Aβ_42_ and increasing concentrations of **MJ040** were added to cells and incubated for 48 h. Addition of Aβ_42_ resulted in a pronounced reduction in cell vitality (~33%, measured as a function of impaired cellular metabolic activity), and the addition of **MJ040** to the extracellular medium was not capable of significantly rescuing the cells (**Fig. 4a**). In a bid to improve cellular activity, the carboxylic acid of the compounds was masked as a methyl ester, to generate the prodrug **MJ040X** (**Fig. 4b, Supplementary Synthetic Procedures**). This compound displayed poor inhibitory activity *in vitro* (**Supplementary Fig. 12**), suggesting the carboxylate is necessary for interaction with the peptide. In the cell-based assays, however, the compound resulted in a significant increase in cell vitality **(Fig. 4a)**. The rescuing effect suggests that the lack of activity observed for the original compound **MJ040** is caused, at least in part, by poor cellular uptake as a consequence of the anionic group. The rationally designed prodrug **MJ040X** successfully permeates into the cells, where the ester is then hydrolysed to generate the free active drug, which inhibits the aggregation process in the cellular environment.

**Figure 4.**
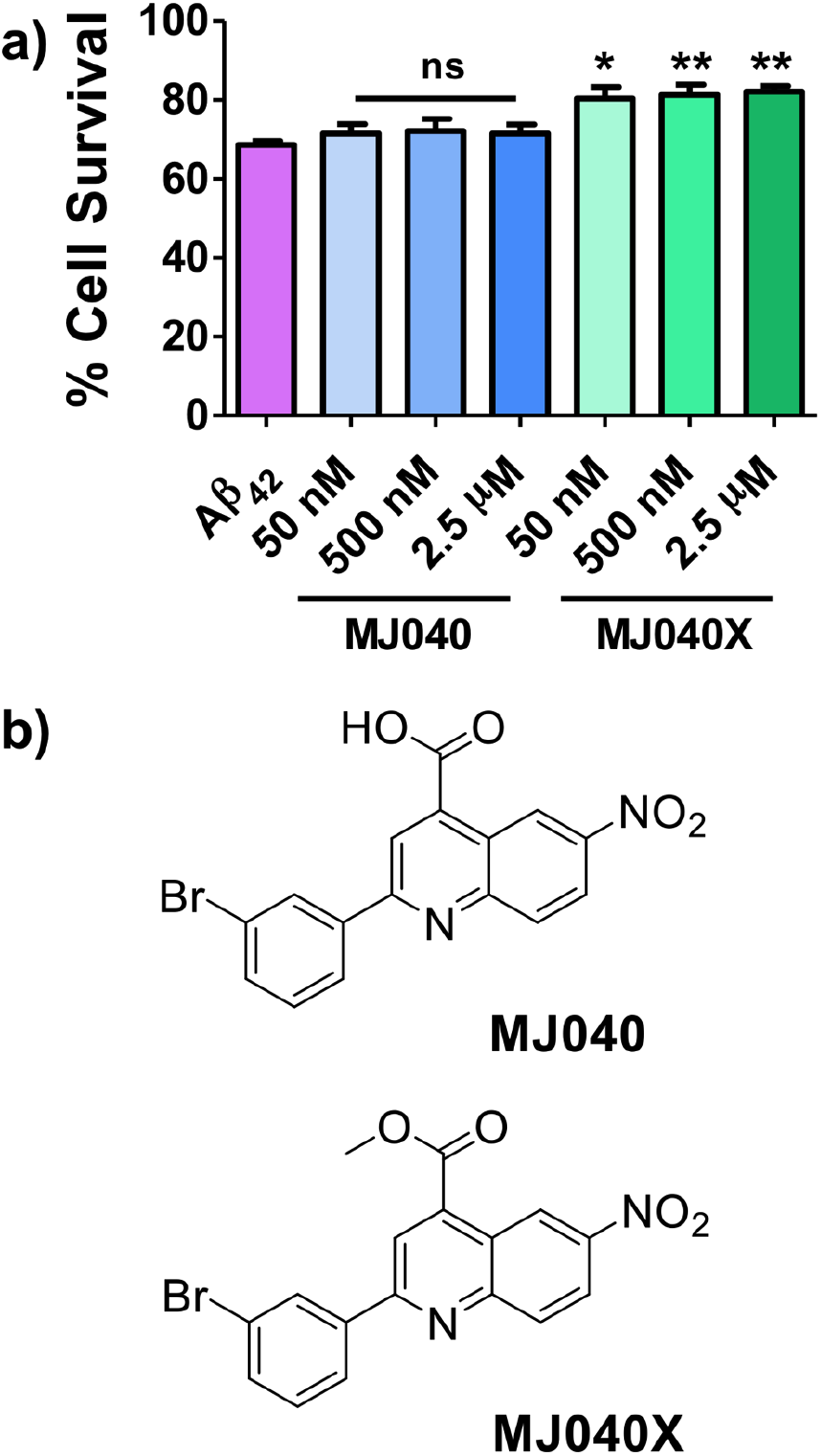
Methyl ester prodrug MJ040X rescues neuronal cells from Aβ_42_-induced cellular stress. **a)** Cell vitality tests monitoring the rescuing effect of **MJ040** and **MJ040X** from monomeric Aβ_42_-induced toxicity. Aβ_42_ monomers (500 nM) with or without the presence of small molecules were added to SH-SY5Y cells. Following 48 h incubation, the cell vitality was assessed using a MTT cell vitality assay. Anionic **MJ040** was unable to rescue Aβ_42_ induced cellular stress, but prodrug **MJ040X** could significantly increase cell vitality. The vitality of untreated cells was set as 100%. Error bars represent SEM, n = 4, statistical analysis performed by one-way ANOVA with Dunnett’s multiple comparison post-test; *p<0.05; **p<0.01. The rescuing effect of **MJ040** and **MJ040X** on preformed Aβ aggregate-induced cell stress is shown in **Supplementary Figure 13**, and the rescuing effect of **MJ001** and **MJ042** in **Supplementary Figure 14b)** Structure of original hit **MJ040** and methyl ester prodrug **MJ040X**.

### Development of a live cell fluorescence lifetime sensor assay confirms cellular Aβ_42_ anti-aggregation activity of hit compound

There is a severe shortage of methods to monitor Aβ_42_ self-assembly in cellular models in real time. Amyloid aggregates in fixed cells can be imaged with immunological staining, which suffers cross reactivity issues with the amyloid precursor protein and its derivatives,^38, 39^ or with the use of amyloid sensitive dyes such as ThT and Congo Red. These effectively stain fixed aggregates, but are generally unsuitable in dynamic live cell studies, as they cannot detect small aggregates low in β sheet content and can induce a mild inhibitory effect on the aggregation process.^40^ Several studies have monitored the uptake of labelled Aβ species, and tracked their movement throughout the cell.^41, 42^ Whilst useful in terms of localisation studies, these cannot report on the underlying aggregation processes and what aggregated species exist at particular time points, as the peptide is uniformly labelled and will exhibit similar fluorescence intensity in a monomeric or aggregated state. Invasive protocols are therefore required to quantify the formation of aggregates. Current cellular methods rely almost exclusively on cell vitality tests. Although these provide a means to quickly elucidate if a compound or the specific Aβ species it generates are toxic to cells, such work does not provide insight into how the compounds are actually acting within the cellular environment, as they do not inform on how the peptide functions within the cells or its aggregation state, and instead only highlight damage to the membrane or dysfunction of the metabolic system.^43, 44^

In order to remedy these shortcomings, we considered that the fluorescence lifetime sensor could be used to follow the uptake and any subsequent aggregation of Aβ_42_ in the presence of hit compounds in SH-SH5Y cells, allowing a direct comparison of Aβ_42_ aggregation propensity *in vitro* and in live cells. The transport of partially labelled Aβ_42_-488 (250 nM) from the extracellular medium into the SH-SY5Y cells was monitored and a drop in the fluorescence lifetime of the attached fluorophore, indicative of peptide aggregation, observed after 12 h (**Fig. 5a**).^25^ To test if the system could be used for small molecule inhibitor screening, the previously reported inhibitor EGCG was employed.^45^ It was found that addition of EGCG to the extracellular medium at the same time as Aβ_42_ addition had little inhibitory effect on intracellular peptide aggregation, with both conditions reaching the same fluorescence lifetime value (**Supplementary Fig. 15**). However, pre-incubating the cells in a drug solution for one hour prior to the addition of the peptide, was shown to significantly inhibit the aggregation in the cells, as measured at 12, 24 and 48 h (**Fig. 5a**). The measurable change in the fluorescence lifetime observed here is directly comparable to that monitored in the *in vitro* format, permitting a comparative analysis of compound activity in both formats, which is not possible by any other method. This protocol was validated by testing a range of other known inhibitory small molecules, which were each seen to inhibit the aggregation to different degrees (**Supplementary Fig. 16**). Treatment with **MJ040** reduced the extent of Aβ_42_ aggregation relative to that of peptide alone, and the modified **MJ040X** provided an even stronger inhibitory effect (**Fig. 5b,d)**. This supports the working idea that **MJ040** displays limited permeability as a result of its anionic centre, and that masking this functionality confers more desirable pharmacokinetic properties. In order to show that the aggregation inhibition did not occur during the pre-incubation of the drug with Aβ_42_ prior to their uptake into cells but indeed intracellularly, we also tested **MJ040X** using a HEK293 cell line overexpressing mCherry-Aβ_42_ intracellularly.^46^ As shown in the **Extended Data Figure 2, MJ40X** was capable of significantly reducing mCherry-Aβ_42_ aggregation as detected by FLIM analysis.

**Figure 5:**
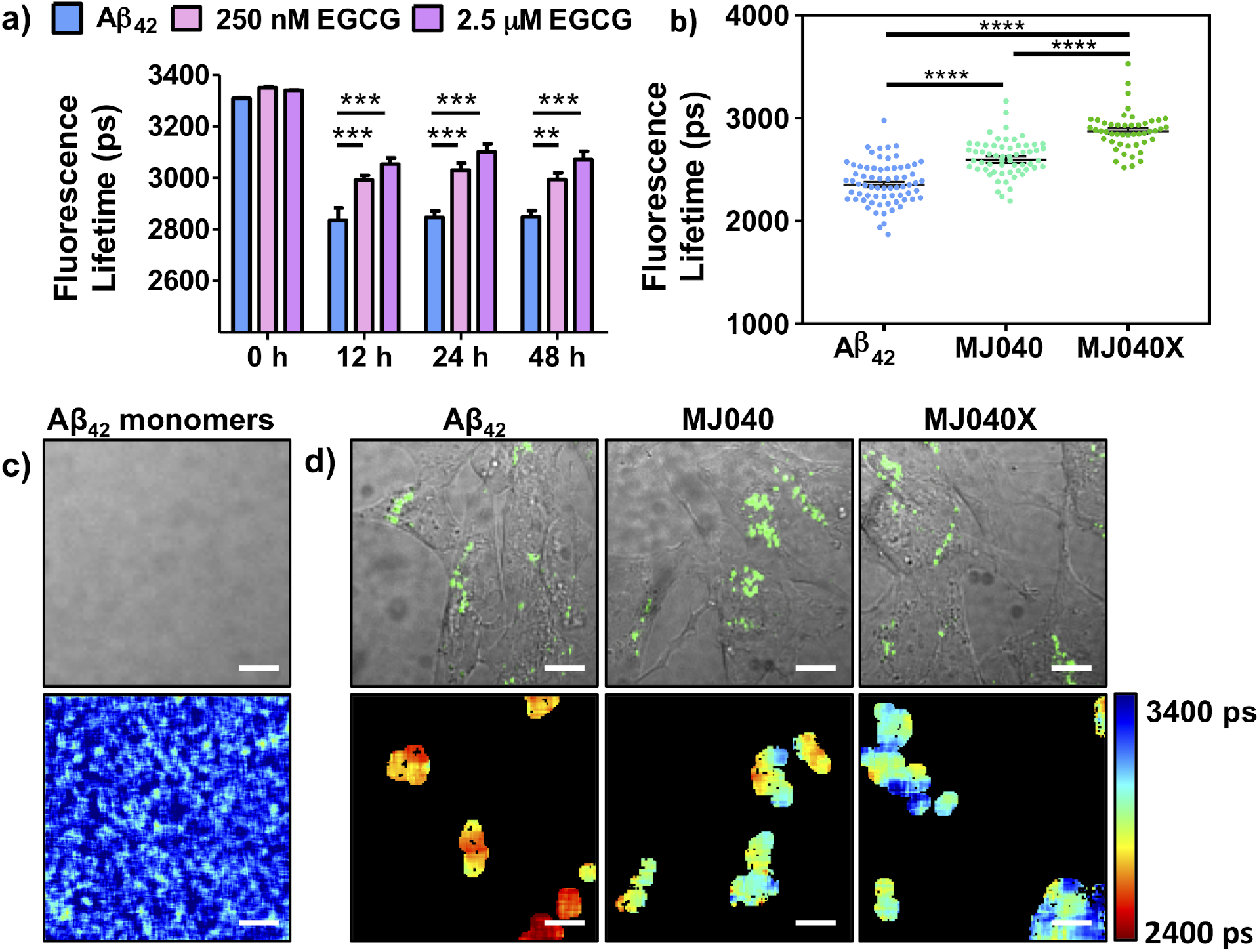
The change in fluorescence lifetime of labelled Aβ_42_ reports on its aggregation in live cells, which can be inhibited with the use of small molecules. **a)** Bar diagram displaying the mean fluorescence lifetime values of partially labelled Aβ_42_ after 12, 24 and 48 h when incubated in the presence of 250 nM or 2.5 μM of EGCG, a known active aggregation inhibitor. Plot shows mean fluorescence lifetime + SEM, statistical analysis performed by two-way ANOVA with Bonferroni post-test, n = 18-30 (see Supplementary Table S2). b) Dot plot displaying the mean fluorescence lifetime values of Aβ_42_-488 obtained following 48 h treatment with either **MJ040** or **MJ040X** (2.5 μM). The prodrug **MJ040X** shows a greater effect on reducing peptide aggregation. The plot shows mean fluorescence lifetime + SEM, statistical analysis was performed by one-way ANOVA with Tukey’s multiple comparisons post-test, based on three biological repeat experiments in which 48-60 cells were analysed (see Supplementary Table S3). **c)** The brightfield/fluorescence and fluorescence lifetime image of 50% labelled Aβ_42_-488 monomers (250 nM) in cell culture medium. Scale bar = 10 μm **d)** Brightfield/fluorescence (above) and corresponding fluorescence lifetime image (below) of SH-SY5Y cells following 48 h incubation with Aβ_42_-488 and hit compounds. Aβ_42_- 488 (250 nM, 50% labelled) with or without the presence of **MJ040** (2.5 μM) or **MJ040X** (2.5 μM) was added to the extracellular medium of the cells. The cells to receive **MJ040** or **MJ040X** treatment were pre-incubated with the compound (2.5 μM) for 1 h prior to Aβ_42_-488 addition. Following 48 h incubation, the cells were washed with medium and the fluorescence lifetime of the internalised Aβ_42_-488 analysed. Scale bar = 20 μm. A typical time trace and exponential decay fits for the data can be found in **Supplementary Figure S17**.

### Hit compound MJ040X inhibits Aβ_42_ aggregation in whole organism disease model

Whole organism studies using a *C. elegans* disease model were carried out to demonstrate the ability of our unified fluorescence lifetime sensor assay to report on aggregation in matched *in vitro* and *in vivo* formats. Worm models have been extensively employed for studying the aggregation of proteins associated with neurodegenerative disease.^47, 48, 49^ Here, a Pmyo3::GFP::Aβ_42_ construct was used (**Supplementary Figure 18)**. In this model system, the myo-3 promoter (Pmyo-3) drives expression of GFP-Aβ_42_. The muscle myosin gene *myo-3* is turned on in post-mitotic embryonic body-wall muscle and, as such, fluorescently labelled Aβ_42_ is easily identified along the periphery of the worms. Similar to the fluorophore previously employed, GFP can act as reporter to inform on the aggregation state of attached amyloidogenic peptides, with the extent of fluorophore quenching indicating the degree of peptide self-assembly.^50^ The fluorescence lifetime of GFP was monitored throughout the life span of adult worms (~15 days), which were either treated with **MJ040X** from the first day of adulthood or directly from the earliest larval stage after synchronisation. A measurable decrease in fluorescence lifetime, indicative of Aβ_42_ aggregation, was observed in accordance with ageing, with a statistically significant reduction observed at day 12 (**Fig. 6a,b,d**). It was found that treatment with the **MJ040X** from adulthood delayed this process. A statistically significant difference in fluorescence lifetime between the treated and untreated control was observed at day 12 of adulthood, but by day 15 peptide aggregation was evident in the treated worms also (**Fig. 6d)**, suggesting a narrow, but significant therapeutic window. Treatment from larval stage, however, prevented Aβ_42_ aggregation until day 15, the last day of measurement for adult worms (**Fig. 6e)**.

**Figure 6:**
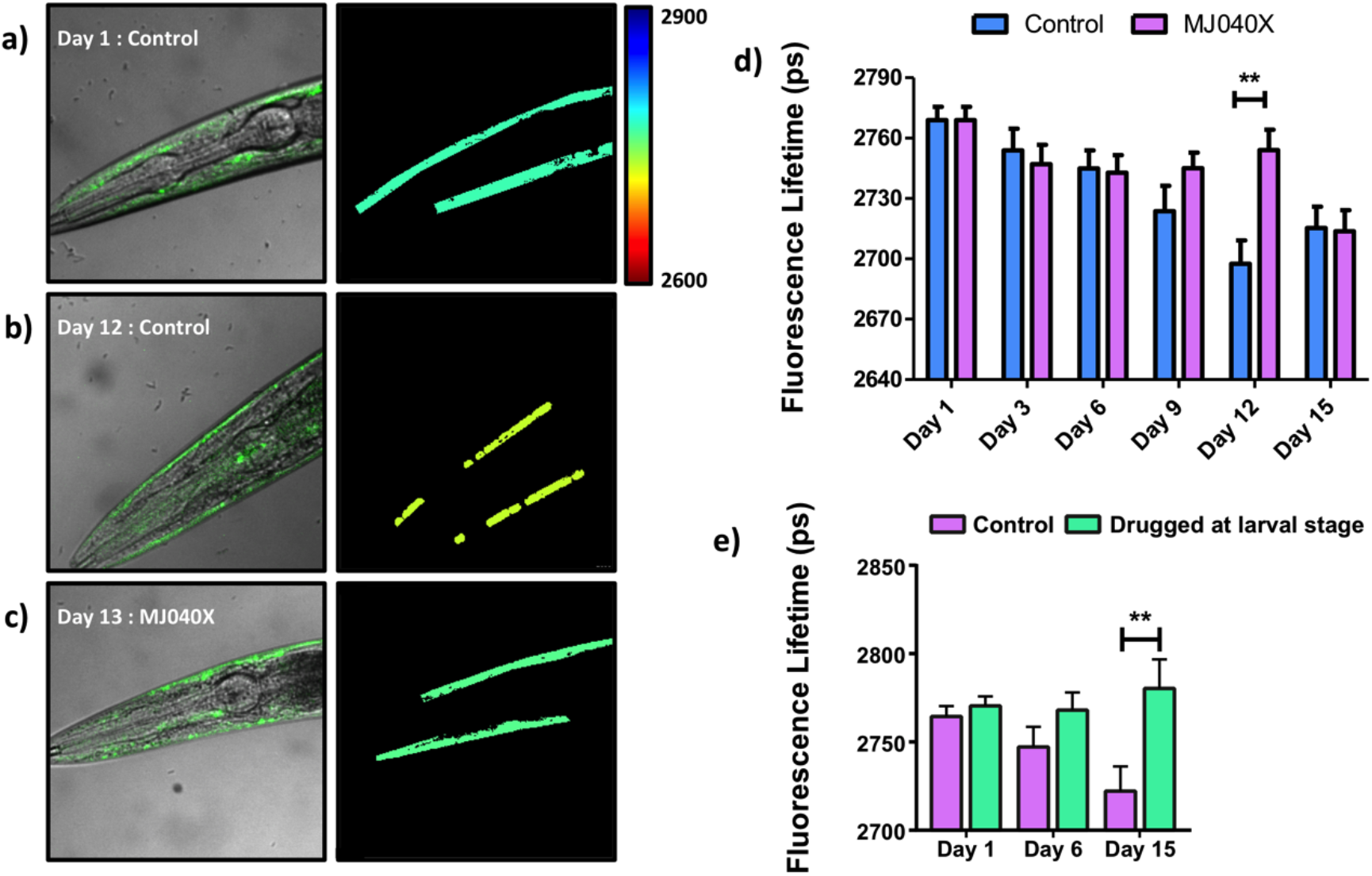
The change in fluorescence lifetime of GFP-Aβ_42_ in disease model *C. elegans* reports on its aggregation in ageing adult worms, and can be used to probe the *in vivo* activity of small molecule inhibitors. Confocal and corresponding fluorescence lifetime images of **a)** day 1 control **b)** day 12 control and **c)** day 12 **MJ040X**-treated worms, expressing GFP-Aβ_42_ in the body muscles. *C. elegans* were grown on OP50 *E. coli* seeded plates, with 0.7 μM **MJ040X**. The worms were changed to new drugged plates on day 8 of adulthood. Scale bar = 20 μm. **d)** Bar diagram displaying the mean fluorescence lifetime values at each time point measured, following the initiation of drug treatment at day 1 of adulthood. A delay in Aβ_42_ aggregation with **MJ040X** treatment was observed in two biological repeats with 8-10 worms analysed for each. Data were analysed by a two-way Anova and a Bonferroni post-test (see Supplementary Table S4). **e)** Bar diagram displaying a comparison of the mean fluorescence lifetime values observed when the *C. elegans* were treated from day 1 of the larval stage. Drug treatment from the larval stage prevents aggregation until day 15, based on two biological repeats with 8-18 worms analysed per repeat. All data are reported as mean fluorescence lifetime + SEM and the statistical analysis was performed using a one-way ANOVA with Sidak’s multiple comparison test (see Supplementary Table S5). A typical time trace and exponential decay fits for the data can be found in Supplementary Figure 17.

## Implications and Conclusions

Recent failures in clinical trials suggest that current aggregation screening strategies are limited in their ability to provide therapeutically viable hit compounds.^12, 19, 51^ Limitations stem from issues in reproducibility (partially due to peptide quality and solubility in a heterogeneous assay),^52^ fluorescence interference caused by intrinsic properties of the test compounds,^35^ and inability to efficiently validate and prioritise hit compounds *in vivo*. There is currently no single method that permits protein aggregation and its inhibition as measured in vitro to be directly correlated to processes in a cellular environment in a mammalian cell or in small organisms. After years of failing to find a suitable therapeutic drug against AD, researchers are still using historically established assays, such as MTT, Thioflavin S staining or fluorescence intensity based aggregate counting to validate their *in vitro* or *in silico* hits.^53, 54, 55^ The unified fluorescence lifetime screening system described in this work overcomes this historical burden and bridges the gap that intrinsically exists between *in vitro* and *in vivo* approaches, permitting results from a medium-throughput microfluidic screen to be directly compared to physiologically relevant cellular and whole organism analysis.

The availability of the nanoFLIM provides a unique opportunity to directly follow the aggregation of Aβ_42_ with large screening libraries, with great sensitivity and temporal resolution, undeterred by limitations that restrict conventional screening approaches. The microfluidic system is advantageous in many respects: (i) *Sample size and reagent consumption*. The nanoFLIM allows for up to 110 replicates per screen, providing an unparalleled sample size and circumventing the issues of irreproducibility that persistently complicate *in vitro* biophysical Aβ_42_ experiments,^12, 23, 56^ even with recombinant material.^29^ The minimised volume (18 nL per droplet vs 100 μL per 96-well plate experiment) necessitates only 219 μg of Aβ_42_ to screen all 445 compounds – with 10 replicates - in nanoFLIM. This is 50-fold less than the 10.7 mg of Aβ_42_ in a ThT experiment performed in 24 96-well plates (see Supplementary Information for details of the comparisons). The miniaturisation and automation reduces time spent performing laborious peptide preparation protocols and makes screening precious novel libraries more feasible. (ii) *Costs*. A 1115-fold reduction in the peptide costs per single experiment is achieved with the nanoFLIM, calculated as 0.087 p per droplet using this assay and £0.97 per well in a 96 well plate format. (iii) *Avoidance of artefacts*. The correct assignment of the inhibitory activity of four spectroscopically active compounds, as validated by TEM imaging analysis, suggests that the fluorescence lifetime sensor is less susceptible to the detection of false positive and negative results than the ThT fluorescence assay. This conventional assay is more prone to bias as a result of intrinsic properties of the screening libraries,^35^ which generally arises from fluorescence interference caused by spectroscopic properties of the test compounds or through competition with the ThT binding site or interaction with the ThT itself.^35, 57, 58^ Removal of these limitations with the nanoFLIM assay increases the true hit rate, and its impact is demonstrated by a screen of DOS- and medicinally-focused compound collections, in which the nanoFLIM permitted the detection of inhibitory activity that was *not* picked up by the conventional ThT assay.

A powerful attribute of the nanoFLIM is the unique fluorescence lifetime readout, which can also be applied for measurements in live cells and *C. elegans* disease models, thereby provided a means to probe amyloid aggregation and the inhibitory effect of added small molecules in multiple formats along the drug development pipeline. The aggregation of Aβ_42_ *in vivo* is considerably more complex than reductionist *in vitro* experiments and remains poorly understood, leading to conflicting interpretations and precluding experimental elaboration of lead compounds that work in an organismal context.^20^ This use of the fluorescence lifetime sensor demonstrates how hit compounds identified in microfluidic droplet screening can quickly and easily be assessed in live cells and disease model whole organisms. Previously, there has been no other single technique that could be used to measure amyloid aggregation in these distinct relevant formats, and this capability gives an inimitable opportunity to reduce attrition rates by bridging *in vitro* and *in vivo* studies. Furthermore, there is great potential to start the compound screening process in the living systems, where the cooperative effects of chaperones, internal surfaces, molecular crowding and other intracellular factors are better taken into consideration. As such, there would be a reduced chance of missing active compounds that do not function in the simplified *in vitro* format, or favouring the detection of non-toxic compounds or those capable of penetrating biological membranes. This system is amenable to a high-throughput scale up through the use of a 96 well plate format of the *in vivo* systems, which could also be easily automated, further speeding up the screening process.

To illustrate the potential of this system, a pilot screening campaign was performed with 445 compounds from medicinally-relevant chemical libraries, yielding a total hit rate of 13% (>30% Aβ_42_ aggregation inhibition). The lead inhibitor identified, **MJ040**, and rationally designed prodrug **MJ040X**, were shown to exert strong inhibitory effects in *vitro*, in live cells and in disease model *C. elegans*, emphasising that biologically active inhibitors can be identified through this comprehensive assay platform. We also believe that it could be easily adapted to screen for aggregation inhibitors of other amyloidogenic proteins, including functional bacterial amyloids, thereby holding potential for the identification of hit compounds for the treatment of a range of amyloid disorders and bacterial pathogenesis.^59^

The breadth of the fluorescence lifetime screening platform and the potential high-throughput afforded by its use, will expedite the rate at which hits are identified, validated *in vivo* and prioritised for future hit development strategies, with the attrition rate in moving through these stages minimised by the unified analysis. The efficiency afforded by this approach has already yielded a lead in **MJ040X**, but the high hit rates observed here suggest further campaigns with existing industrial compound libraries are likely to identify further modulators of the aggregation of Aβ_42_, providing new opportunities to overcome the current unsatisfactory situation, where no Alzheimer’s disease treatments exist.

## Methods

### Reagents

Synthetic Aβ_42_ (>95%, Eurogenentec) and Aβ_42_ Hilyte™ Fluor 488 (>95%, labelled at the N-terminus) were purchased from Anaspec as lyophilised powder. ThioflavinT was purchased from AbCam. The compound screening libraries were obtained from the Spring Research group (Department of Chemistry, University of Cambridge). All other chemicals, unless otherwise stated, were purchased from Sigma Aldrich.

### Peptide Preparation

The peptide was prepared as previously described.^30^ Briefly, lyophilised Aβ_42_ (1 mg) was dissolved in ice cold trifluroacetic acid (200 mL), sonicated at 0 °C for 60 s, then lyophilised overnight. Ice cold 1,1,1,3,3,3-hexafluro-2-propanol (1 mL) was added, sonicated at 0 °C for 60 s and aliquoted into 20 μL portions. The samples were lyophilised overnight and were stored at –80 °C until use. The concentration of the aliquots was determined using amino acid mass spectrometry analysis. The required concentration of unlabelled Aβ_42_ was prepared by dissolving the solution in dimethyl sulfoxide (DMSO) (5% of total solvent volume), then adding sodium phosphate buffer (NaPi, 50 mM, pH 7.4). Prior to use, the solution was sonicated at 0 °C for 3 min, centrifuged at 13,400 rpm at 0 °C for 30 min to remove preformed aggregates. Lyophilised Aβ_42_ Hilyte™ Fluor 488 peptide (0.1 mg) was dissolved in 1% NH_4_OH (200 μL) and sonicated for 60 s at 0 °C. The sample was aliquoted into 5 μL units, snap frozen in liquid N_2,_ then stored at –80 °C. Before use, the sample was thawed on ice and NaPi buffer was added to bring the solution to the required concentration.

For studies with partially labelled peptide, each peptide was prepared as above then mixed at the appropriate ratios before each set of experiments. This was aliquoted into small units, then snap frozen and stored at –80 °C until use.

### Microfluidic Device Fabrication

The silicon master mould was fabricated by MicroLiquid (Gipuzkoa, Spain) using a two layer soft lithographic technique, as previously described.^60, 61^ The depth of the first layer with a serpentine channel was 175 μm, while the depth of the square traps were 250 μm. PDMS replicates of these devices were bonded to a thin glass coverslip (thickness 130 μm) using oxygen plasma. The devices were silanized by pipetting a fresh solution of 2% Trichloro(1H,1H,2H,2H-perfluorooctyl)silane in HFE-7500 (3M) into the chips. Subsequently, PTFE tubing (diameter: 200 μm) was manually inserted in designed entrance channels and glued in place by curing PDMS over. This ensured air-tight connection as well as preserving the order of the droplets as they transited from tubing to chip. One tubing was then connected to a syringe pump (Chemyx Fusion 200) operating in withdrawal mode, while the other tubing was inserted and clamped into the stainless steel hook of a Mitos Dropix. A gas-tight glass syringe (100 μL) was used to fill the device with HFE-7500 oil containing 1% Pico-Surf surfactant (Dolomite Microfluidics) and the chip was inspected to confirm the absence of air bubbles. Next, 10 μL of each compound was pipetted into the loading strip of the Dropix. The droplet sequence was programmed to obtain 18 nL droplets with 36 nL oil spacing between each drop. Typical flow rate for producing the droplets was 2 μL/min. Before reaching the device, the droplets were slowed down to 1 μL/min and the filling process was monitored with a bright-field camera of an inverted microscope (Olympus, IX71). After completion of the filling process, the flow was stopped, unless specified in shearing experiments.

### Spectroscopic Assays

Final concentrations of 20 μM ThT with 10 μM unlabelled peptide in sodium phosphate (NaPi, 50 mM) were used for all samples. For screening the chemical libraries, 50 μM of the test compound was added. The assay samples (total volume 100 μL) were mixed in a black non-binding 96-well plate (Greiner Bio-One, Switzerland). The plate was sealed (Nunc™, polyolefin acrylate film) and loaded into the fluorescence plate reader (Tecan, Switzerland) at 37 °C. Fluorescence kinetics were measured at 5 min reading intervals, with 15 sec shaking before each read. The excitation and emission wavelengths were 440 nm and 480 nm, respectively.

### Fluorescence lifetime imaging microscopy

Fluorescence lifetime imaging was performed on a custom-built confocal microscope (FV300, Olympus) as previously described^21^ that utilises a time-correlated single photon counting (TCSPC) module (SPC-830, Becker & Hickl GmbH) and is shown in the Extended Data Figure 3 and 4, Supplementary Figure S17). Briefly, a supercontinuum source (SC390, Fianium) operating at a 40 MHz repetition rate was used for excitation. The excitation light was filtered using an acousto-optic tunable filter (AOTFnC-400.650, QuantaTech) centered at 480 nm to excite GFP and AF488 or at 585nm to excite mCherry. Fluorescence emission from the sample passed through a band-pass filter (FF01-525/39-25 or FF01-624/40-25, Semrock) before reaching the detector (PMC-100, Becker & Hickl GmbH). The photon detection rate for each pixel was kept below 1% of the laser repetition rate in order to avoid photon pile-up. Air objectives (PlanApo 2x and 40x, Olympus) were used for imaging the microfluidic chip and *C. elegans*, respectively. An oil objective (PlanApo 60x BFP1 C2, Olympus) was used to image cells. All FLIM data were analysed using either commercial software SPCImage (Becker & Hickle GmbH) or open-source FLIM-fit software.^62^ A biexponential fit was used for nanoFLIM and for FLIM data from AF488 containing cells. A single exponential fit was used for FLIM data from *C elegans* data and from mCherry containing cells (Extended Data Figure 2). The phasor plot analysis^62^ feature in SPCImage was used to validate the use of a biexponential fit on certain FLIM data (Extended Data Figure 4).

### AFM

A freshly cleaved mica surface was prepared by sequential treatment with potassium hydroxide and 0.1% poly-lysine solution. After the specified time of incubation (20 μM Aβ_42_, 100 μM compound), samples were transferred directly to the slides and were allowed to dry for 30mins. Samples were then rinsed with Milli-Q water and dried in the air. AFM images were acquired on a commercial system (Bioscope RESOLVE, Bruker) and Nanoscope software (Bruker). The instrument was operated in tapping mode in air.

### Transmission Electron Microscopy (TEM)

Samples were stained with 2 % (w/v) uranyl acetate. Images were obtained at various magnifications using a Tecnai G2 80-200kv transmission electron microscope transmission electron microscope and captured with a bottom mounted digital camera (AMT).

### Dot blot

20 μM Aβ_42_ and 100 μM compound were incubated at 37 °C. At specified time points, 10 μL aliquots were removed and stored at −20 °C until use. 5 μL samples were spotted onto a nitrocellulose membrane (Amersham Hybond ECL, GE Healthcare Life Sciences) and were allowed to dry for 1 h. The membranes were blocked with 5% non-fat milk in tris-buffered saline (TBS), then washed with TBS. The membranes were treated with the primary antibodies A11 (Invitrogen) and 6E10 (Invitrogen) for 12 h. After washing (0.01% Tween20 in TBS), anti-rabbit horseradish peroxidase (HRP) conjugated antibodies (ThermoScientific) were added. After 2 h, the membranes were washed (0.01% Tween20 in TBS). An enhanced chemiluminescent (ECL, Thermo Scientific Pierce™) substrate was added and the samples exposed to X-ray film, as per manufacturer instructions. Positive A11 stained dots are indicative of the presence of oligomeric species, but not Aβ monomer or fibrils. 6E10 stained dots are indicative of any Aβ species, irrespective of the conformation.

### SH-SY5Y cell culture

Human neuroblastoma cells (Sigma-Aldrich, Gillingham, UK) were grown in a serum-containing medium (SCM) consisting of 15% FBS, 1% non-essential amino acids (Sigma), 1% L-glutamine (Life Technologies, UK) and 1:1 minimal essential medium (Sigma) and nutrient mixture F-12 Ham (Sigma). For cell viability assays and uptake experiments the cells were cultured using a serum free medium (SFM), in which the FBS was replaced with 2% B27 complement (Life Technologies).

### mCherry HEK cell culture

Flp-InTM T-RExTM 293 cell line (Invitrogen), with a stably integrated FRT site and a TetR repressor, served to create new stable cell lines expressing mCherry-Aβ42(WT), i.e. pcDNA5/FRT/TO-mCherry-Aβ(WT) under the Flp InTM expression vector.^46, 63^ All cell lines were grown in cell culture medium containing: Dulbecco’s Modified Eagle’s Medium (DMEM), fetal bovine serum (FBS; 10%; Thermo Fisher Scientific), antibiotic-antimycotic (1%; Thermo Fisher Scientific), and glutaMAX (1%; Thermo Fisher Scientific). Cells were incubated in T75 flasks at 37 °C, in a 5% CO_2_ atmosphere, and passaged approximately every 3–4 days to reach 80–90% confluency. Cell lines were regularly tested for mycoplasma contamination with the MycoAlertTM PLUS Mycoplasma Detection Kit (Lonza, Walkersville).

### Cell Vitality Assay

Monomeric peptide experiments: SCM containing the test compounds were added to the SH-SY5Y and were incubated for 1 h. The media was removed and the cells were rinsed with minimum essential medium (MEM). Aβ_42_ monomers (500 nM) with or without the presence of test compound in SFM were added. Following 48h incubation the cytotoxicity was assessed using an MTT cell viability assay (measuring cellular metabolic activity as an indicator of cell vitality), according to the manufacturer’s instructions (Invitrogen). Briefly, medium was aspirated from the cells and replaced with 100 μL of fresh medium and 10 μL of 12 mM MTT. After incubation for 4 h at 37 °C, 85 μL of the medium was removed and 50 μL of DMSO added. The samples were mixed thoroughly, incubated overnight at 37 °C for 10 min, then the absorbance was recorded at 540 nm.

Pre-aggregated peptide experiments: Aβ_42_ (10 μM) was pre-incubated for 24 h with or without the test compounds. The sample were diluted with SFM (500 nM Aβ_42_) and added to SH-SY5Y cells. After 48 h treatment, cell vitality was evaluated using an MTT assay as previously described.

### Cellular fluorescence lifetime sensor experiments

SH-SH5Y cells (30,000 cells/well) were plated in Lab-Tek II chambered coverglass plates (NUNC™, Thermo Fisher Scientific, Cramlington, UK) in SCM, and were incubated for 24 h. The media was removed and fresh SCM containing the test compound of interest, or vehicle control, was added. The cells were allowed to incubate at 37 °C for 1 h. The media was removed and SFM containing 50% labelled 250nM Aβ_42_-488 and the test compound was added to the cells, which were then incubated for 24 or 48 h. mCherry-Aβ42(WT) cells (20,000 cells/well) were plated in Lab-Tek II plates (Nunc™, Thermo Fisher Scientific) containing cell culture medium supplemented with tetracycline (1 μg/mL), to induce construct expression. The compound of interest (M040X) or vehicle control (DMSO) was added. MJ040X was first dissolved in DMSO to make a stock solution (20 mM), and then further diluted withNaPi buffer (50 mM, pH 7.4) to make a working solution (1 mM), which was then added to cell culture medium to obtain the required final concentration (2.5 μM). Samples were incubated for 48 h (at 37 °C, 5% CO_2_) before analysis. The cells were rinsed 3 times with MEM then imaged in a chamber at 37 °C and 5% CO2 on the microscope stage. 8-10 images were taken per condition, with 3-10 cells per image, depending on magnification. The FLIM data was analysed using the FLIMfit software, using image-wise analysis on segmented regions.^64^

### Worm model experiments. *C. elegans*

(Pmyo3:: GFP::Aβ42; expressing Aβ_42_ in its muscle myosin; Fig. S21) were cultivated on a diet of OP50 *E. coli* and were maintained at 20 °C on nematode growth medium (NGM) according to well described protocols.^65^ Compound treatment was achieved by adding solutions of the compound or vehicle control (700 nM, 0.05% DMSO) to the bacterially-lawned agar plates and allowing the solvent to dry off before adding the worms. Worms treated from adulthood were transferred to drugged 5-fluoro-2′-deoxyuridine (FUDR, 75 μM) containing NGM plates 2.5 days post synchronization (NaOH 0.25 M, NaOCl 0.8%, 6 min). Worms treated from larvae were transferred to drugged NGM plates directly after synchronization, then transferred to drugged FUDR plates after 2.5 day. All plates were seeded with concentrated OP50 *E. coli*. Adults were transferred to freshly drugged FUDR NGM plates after 8 days. Imaging was carried out at day 3,6,9,12 and 15. Two repeats of each time point were taken, with 8-10 worms imaged per condition for day 1 adult treated worms. Two repeats of each time point, with 8-18 worms imaged per condition for worms treated from larvae.

### Statistical Analysis

Data were analysed using GraphPad Prism software, version 5. All comparisons were accompanied by one-way or two-way analysis of variance (ANOVA), with a Dunnett’s, Sidak or Bonferroni post-test respectively utilised to analyse variance between each set of treatments (see Supplementary Tables S2-4). P-values are annotated * for p<0.05, ** for p<0.01, *** for p<0.001 and **** for p<0.0001.

### Nuclear Magnetic Resonance

To probe the binding properties of MJ040 with Aβ42, we used ^1^H-^15^N HSQC experiments in solution NMR spectroscopy. Recombinant ^15^N-labelled Aβ42 peptides with ammonium acetate counterions were obtained from AlexoTech (Umea, Sweden) and handled on ice at all times. The peptide powder was solubilized in 10 mM NaOH at concentrations of 5 mg/mL and stored at −80 °C until required. To prepare NMR samples, Aβ42 stock solutions were diluted into sodium phosphate buffer (50 mM Na2HPO4/ NaH2PO4) to give a final peptide concentration of 100 μM and a pH of 7.5. NMR experiments were carried out at 5°C on an NMR spectrometer operating at the ^1^H frequency of 700 MHz and equipped with triple resonance HCN cryo-probe. The ^1^H-^15^N HSQC experiments were recorded using a data matrix consisting of 2048 (t_2_, ^1^H) × 140 (t_1_, ^15^N) complex points. Assignments of the resonances in ^1^H - ^15^N - HSQC spectra of Abeta42 were derived from our previous study.^66^

### Modelling

We utilised a set of structures of the Aβ42 peptide selected from clusters of conformations within an ensemble generated using molecular dynamics simulations. Using these clusters, we identified the binding ‘hot spots’ for MJ040 on the monomeric form of the Aβ42 peptide following our published protocol^67^ based on the application of the FRED program.^68^ In order to inform the selection of bound states from the docked structures, we utilised the experimental observations providing the probability of contacts between MJ040 and Abeta42, as derived from the NMR measurements. The results showed that transient hydrophobic clefts are formed in the C-terminal region of the peptide, providing hot-spots for the binding of the hydrophobic parts of MJ040, with hydrophilic groups of the molecule pointing toward the solvent.

## Data Availability

The datasets generated during and/or analysed during the current study are available from the corresponding author on reasonable request.

## Acknowledgements

The authors would like to acknowledge Dr Kaveh Ashrafi and Jason Liu for generating the *C. elegans* strain, Anna Lena Ellermann for help with preparation of figures, Dr Tessa Sinnige for help with *C. elegans* cultures, and thank Dr Della David and Prof. Michele Vendruscolo for helpful discussions. S.C. acknowledges a studentship from the Biotechnology and Biological Sciences Research Council (BBSRC) and support from the Cambridge Trust. FH holds H2020 ERC Advanced (695669) and PoC (665631) awards. G.S.K.S. acknowledges funding from the Wellcome Trust (065807/Z/01/Z) (203249/Z/16/Z), the UK Medical Research Council (MRC) (MR/K02292X/1), Alzheimer Research UK (ARUK) (ARUK-PG013-14), the Michael J Fox Foundation (16238) and Infinitus China Ltd.

## Author contributions

S.C., F.G., L.v.V., D.R.S., F.H. and G.S.K.S. designed research; S.C., F.G., M.J., L.v.V., S.W.-V., C.P., G.F. and A.D.S. performed research; S.C., L.v.V., F.G., C.M., D.D, C.F.K., D.R.S, F. H. and G.S.K.S analysed data; and S.C., L.v.V., D.R.S, F.H. and G.S.K.S wrote the paper with comments and input from all authors.

## Additional information

**Supplementary information** The online version contains supplementary material available at https://doi.org/… and supplementary videos 1-3 showing details of device operation.

**Peer review information** *Nature Chemistry* thanks the anonymous reviewers for their contribution to the peer review of this work.

**Reprints and permissions information** are available at www.nature.com/reprints.

---

**Extended Data Figure 1:**
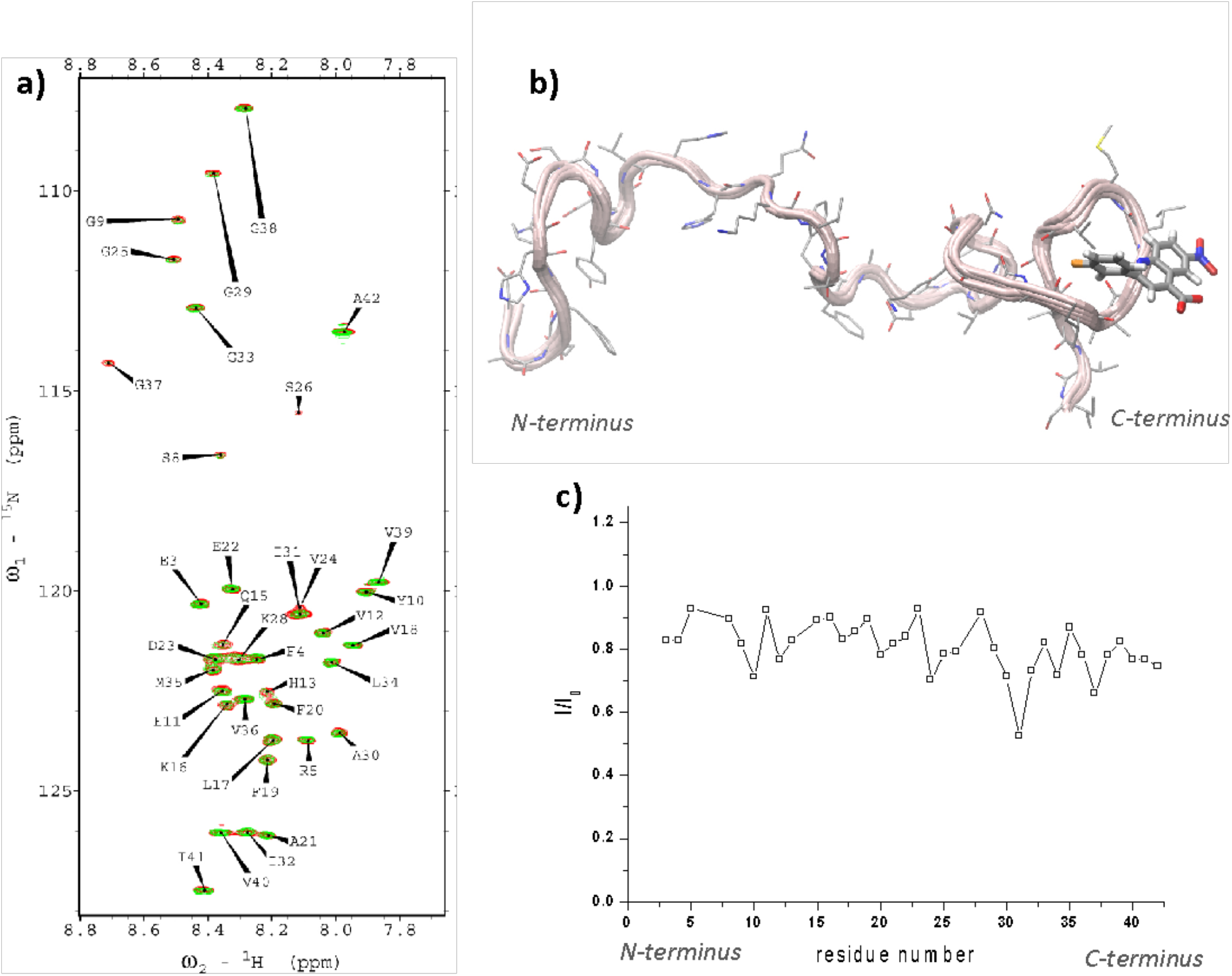
NMR analysis and modelling reveal the binding site of MJ040 in the C-terminal of monomeric **Aβ**_**42**_. a) ^1^H-^15^N-HSQC spectrum of Aβ_42_ (100 μM) measured (50 mM phosphate buffer, pH 7.5, 2% DMSO) in the presence (green) or absence (red) of MJ040X (500 μM). Both experimental conditions included 2% DMSO. The spectra were measured at 278K using an NMR spectrometer operating at the ^1^H frequency of 700 MHz. No significant chemical shift changes were observed upon addition of MJ040X to Aβ_42_. However, the interaction with the molecule induced *broadening* of the resonances, with strongest effects in I/I_o_ found in the C-terminal region (specifically in residues E31 and G37). Assignments of the resonances in ^1^H - ^15^N - HSQC spectra of Aβ_42_ were derived from a previous study.^69^ b) Modelling of the interaction between MJ040X and Aβ_42_ conformations using an approach to identify the binding modes of small molecules with this protein^65^. The programme FRED^68^ was used to perform the docking of the molecule in these hot spots. c) Variation of signal intensity over the sequence of Aβ_42_ in the presence of MJ040X (intensity of the green resonances (with MJ040X)/intensity of the background Aβ_42_ resonance in a).

**Extended Data Figure 2:**
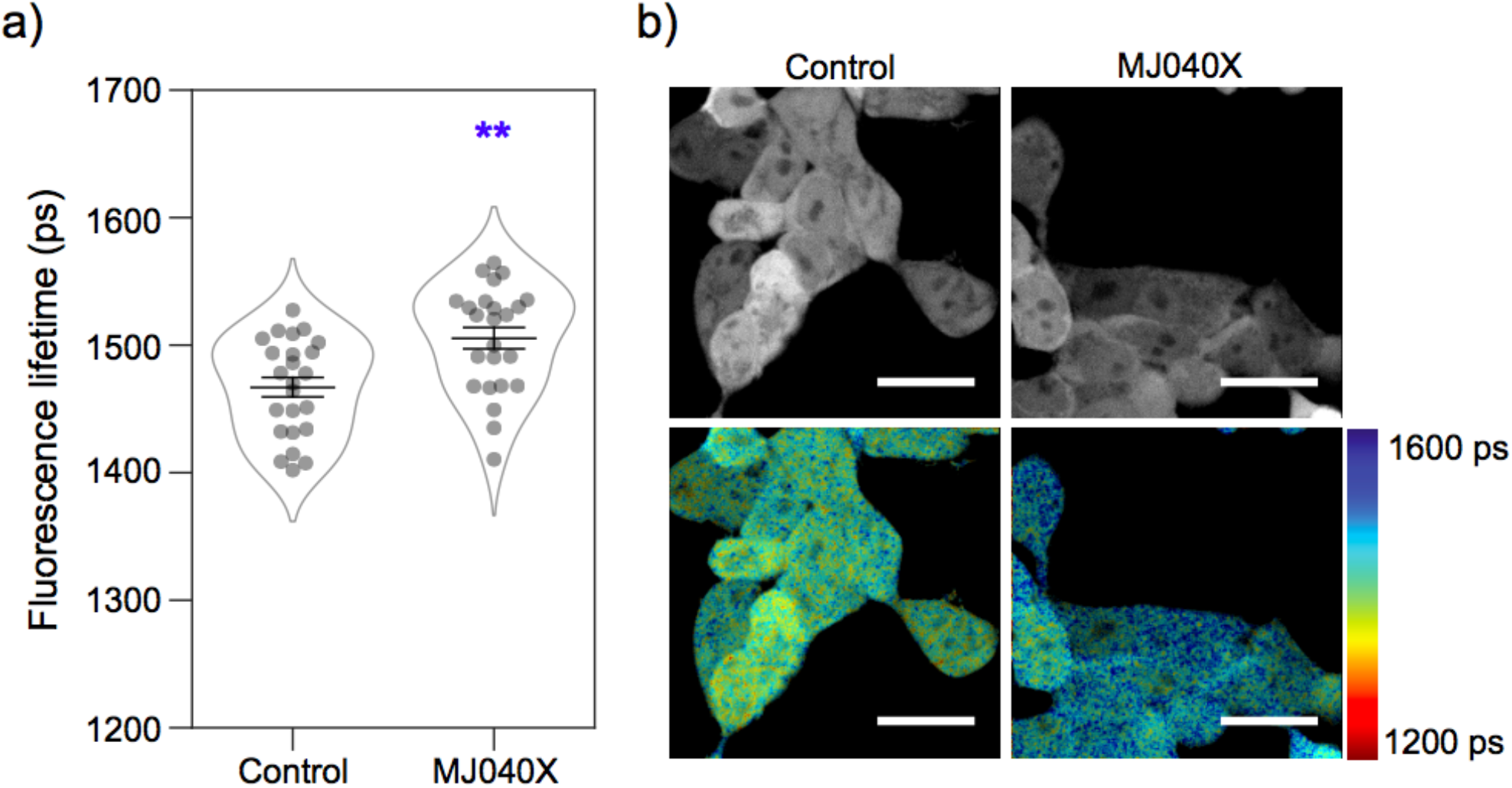
MJ040X treatment increases fluorescence lifetime of mCherry-Aβ_42_ HEK cells. Intensity-averaged fluorescence lifetimes of mCherry-Aβ_42_ cells treated with DMSO (control) or MJ040X (2.5 μM) for 48 h. The intensity-averaged fluorescence lifetimes are 1467 ± 8 ps and 1505 ± 8 ps, for the control and MJ040X-treated cells, respectively, confirming that MJ040X does inhibit the aggregation of Aβ_42_ inside the cell. The diagram displays mean fluorescence lifetime ± SEM. Statistical analysis was performed using an unpaired t-test, 3 biological repeats ** = p<0.01. Scale bar = 20 μm.

**Extended Data Figure 3:**
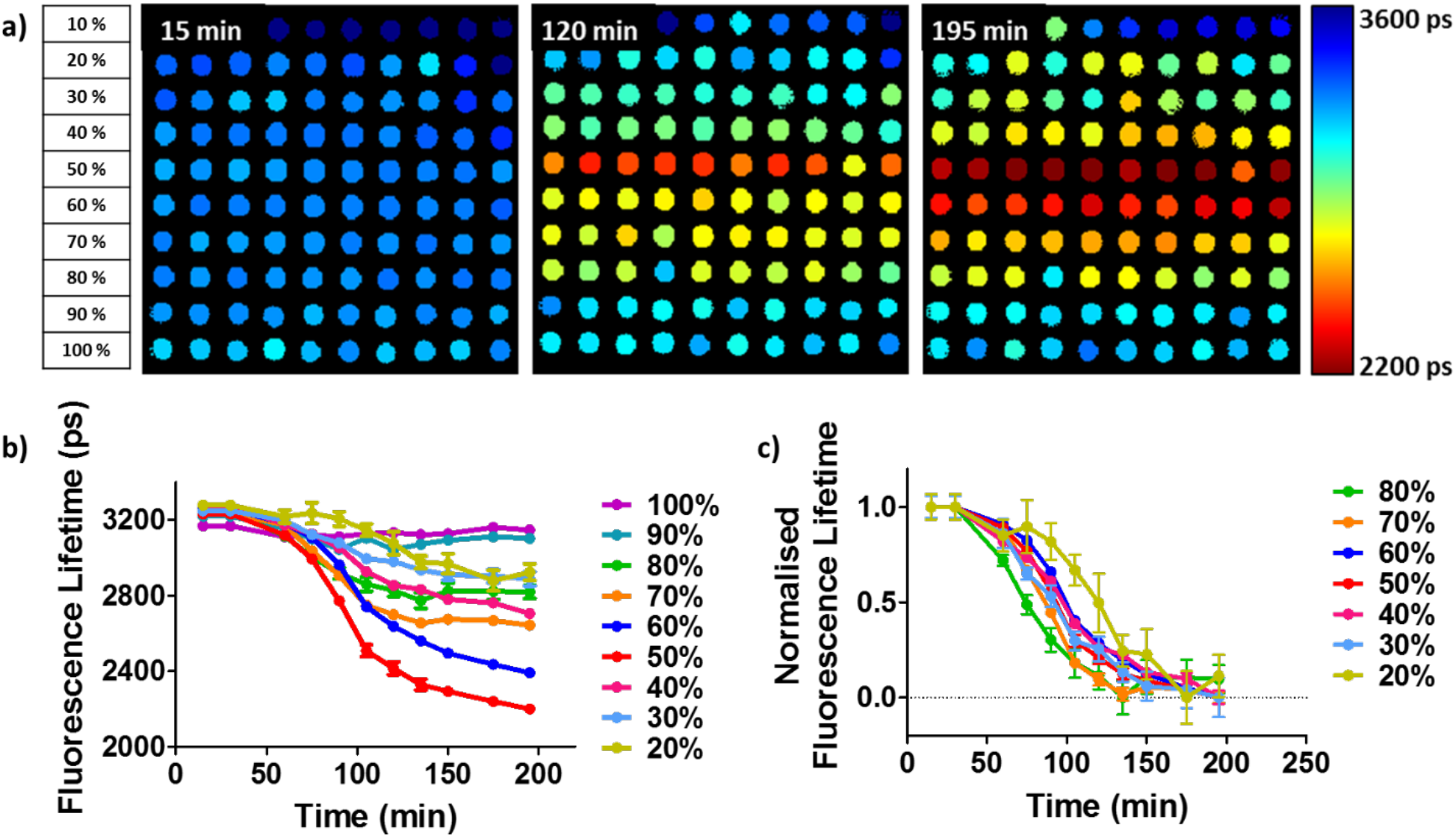
Effect of labelling density on rate of peptide aggregation and change in fluorescence lifetime. **a)** Labelling density gradient in microfluidic chip. Each row contains 10 droplets of increasing labelled Aβ_42_ (10 μM**). b)** Aggregation profiles of Aβ_42_ at different labelling densities. 50% labelled shows the largest dynamic range. **c)** Normalised aggregation profiles at different labelling densities. Aggregation curves from 90% and 100% labelled Aβ_42_ are omitted due to negligible aggregation. Plots show mean ± SEM, n = 10.

**Extended Data Figure 4:**
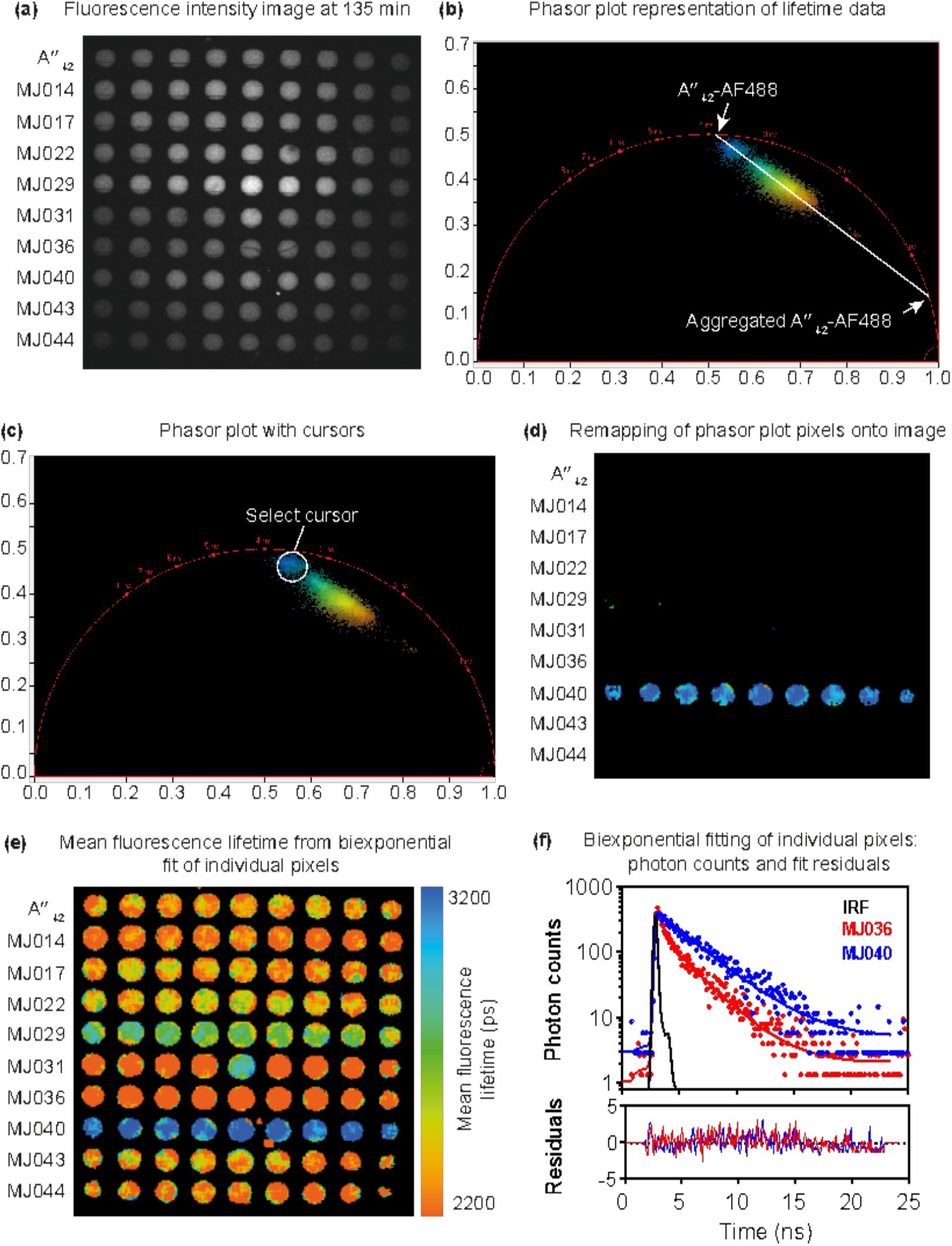
Detailed analysis of compound collection screening with a phasor plot and biexponential fit. (a) Raw fluorescence intensity image of nine droplet replicates containing peptide and test compounds at the 135-minute time point. (b) Phasor plot analysis of the lifetime data from the entire nanoFLIM image. This is an unbiased analysis approach, which does not rely on a fitting model of exponential decay data. Native unaggregated Aβ_42_ tagged with AF488 lies in the universal circle near 4 ns and fully aggregated Aβ_42_-AF488 has a very short (quenched) fluorescence lifetime close to 1 ns. All droplets contain a mixture of these two species in different proportions. (c) The phasor plot shows a bimodal distribution in which a small subgroup of points (selected with the cursor) are closer to the native, unaggregated Aβ_42_ species than the rest of the points. (d) Points selected by the cursor in (c) with the least amount of aggregated Aβ_42_ correspond to the droplets containing MJ040, showing the effectiveness of the compound in inhibiting aggregation. (e) The phasor plot reveals that decay curves in image (a) are biexponential as they lie within the universal circle between two pure single exponential species. Standard biexponential fitting to decay curves are performed to extract quantitative lifetime parameters. The mean fluorescence lifetime is high for droplets containing MJ040, which shows that Aβ_42_ aggregation is inhibited in these droplets. (f) Typical fluorescence decay curves (with a comparison between MJ036 and MJ040), biexponential fits to the raw data, and the fit residuals for individual pixels of the FLIM image. The instrument response function (IRF) for the setup is also plotted.

