## Supplementary Information for "A unified *in vitro* to *in vivo* fluorescence lifetime screening platform yields amyloid β aggregation inhibitors"

Figure S4: Detailed analysis of a collection of known inhibitors in the NanoFLIM assay and comparison to other widely used assays.......................... 6-8

**Supplementary Videos**

Still images from each video are placed below.

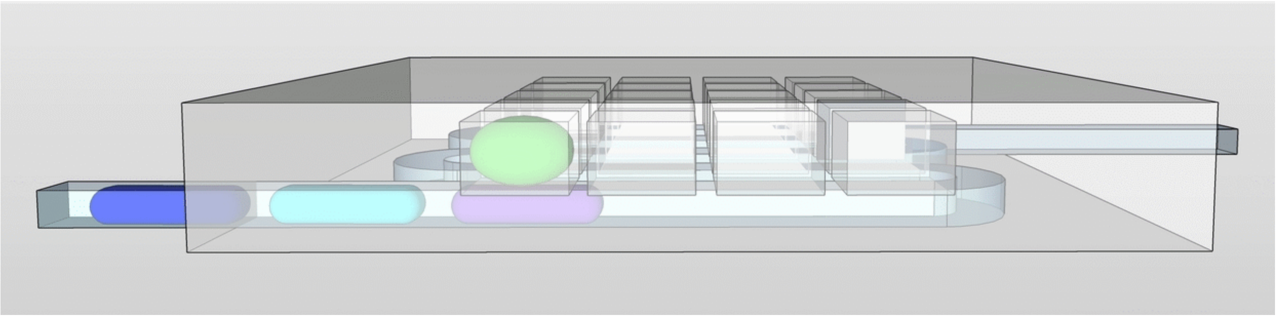

**Video S1:** GIF animations of microfluidic device filling

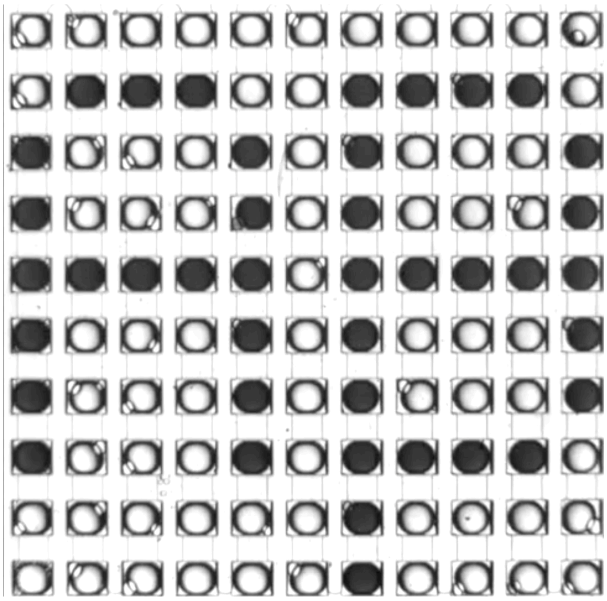

**Video S2:** Filling microfluidic device with predefined droplet sequence

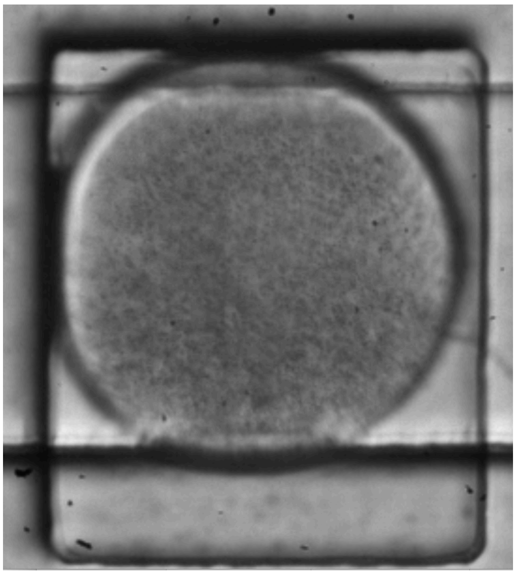

**Video S3:** Inducing a shear force in the device by flowing oil through the channel under the droplets.

### Supplementary Notes

**Supplementary Note 1:** NMR Spectroscopy

For $^{1}H$-$^{15}N$ HSQC experiments in solution NMR spectroscopy, recombinant $^{15}N$-labelled $A\beta_{42}$ peptides with ammonium acetate counterions were obtained from AlexoTech (Umea, Sweden) and handled on ice throughout the experiment. The peptide powder was solubilized in 10 mM NaOH at concentrations of 5 mg/mL and stored at −80 °C until required. To prepare NMR samples, $A\beta_{42}$ stock solutions were diluted into sodium phosphate buffer (50 mM $Na_2HPO_4$/ $NaH_2PO_4$) to give a final peptide concentration of 100 μM and a pH of 7.5. NMR experiments were carried out at 5°C in an NMR spectrometer operating at the $^{1}H$ frequency of 700 MHz and equipped with a triple resonance HCN cryo-probe. The $^{1}H$-$^{15}N$ HSQC experiments were recorded using a data matrix consisting of 2048 ($t_2$, $^{1}H$) × 140 ($t_1$, $^{15}N$) complex points. Assignments of the resonances in $^{1}H$-$^{15}N$-HSQC spectra of $A\beta_{42}$ were derived based on published data.[1]

**Supplementary Note 2:** Modelling

A set of structures of the $A\beta_{42}$ peptide was selected from clusters of conformations within an ensemble previously generated using molecular dynamics simulations.[2]. Using these representative conformations, we identified the binding 'hotspots' for MJ040 in the $A\beta_{42}$ peptide following a published protocol for structural interpretation.[2] NMR measurements provide the probability of contacts between MJ040 and $A\beta_{42}$ and these data were used to further define the hotspots. Hydrophobic clefts, formed in the C-terminal region of $A\beta_{42}$ for some representative conformations of the ensemble, were identified and suggested a binding pocket for the hydrophobic regions of MJ040X, with hydrophilic groups of the molecule pointing toward the solvent. Then the programme FRED[3] was used to perform the docking of the molecule in these hotspots. The interpretation was based on the assumption that the MJ040X initially binds the monomer, but higher order effects cannot be excluded.

### Supplementary Figures

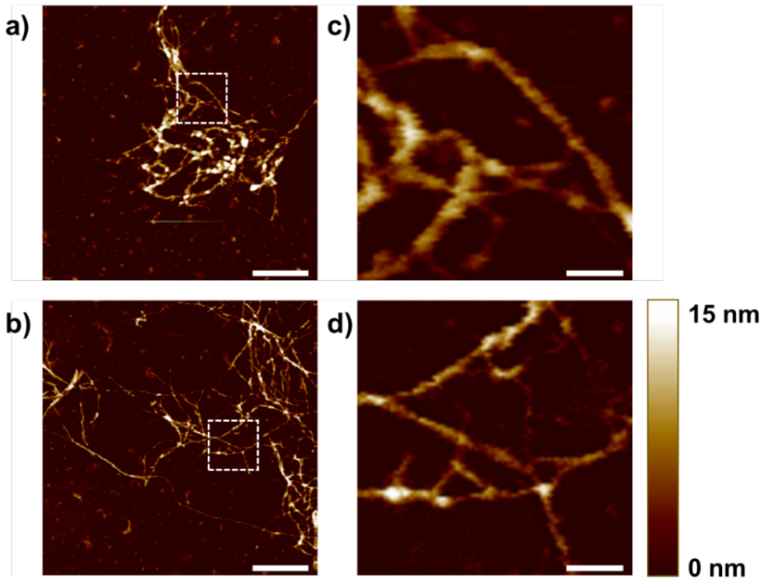

**Figure S1: Effect of 50% labelling density on fibril morphology of Aβ42 monitored by AFM. a)** and **b**) Unlabelled and 50% Hilyte Fluor488 labelled $A\beta_{42}$, respectively, following 7 days of incubation at 37 °C. Scale bar = 600 nm. **c)** and **d)** Zoomed in images of unlabelled and labelled $A\beta_{42}$ showing similarities in fine fibrillar structure. Scale bar = 100 nm.

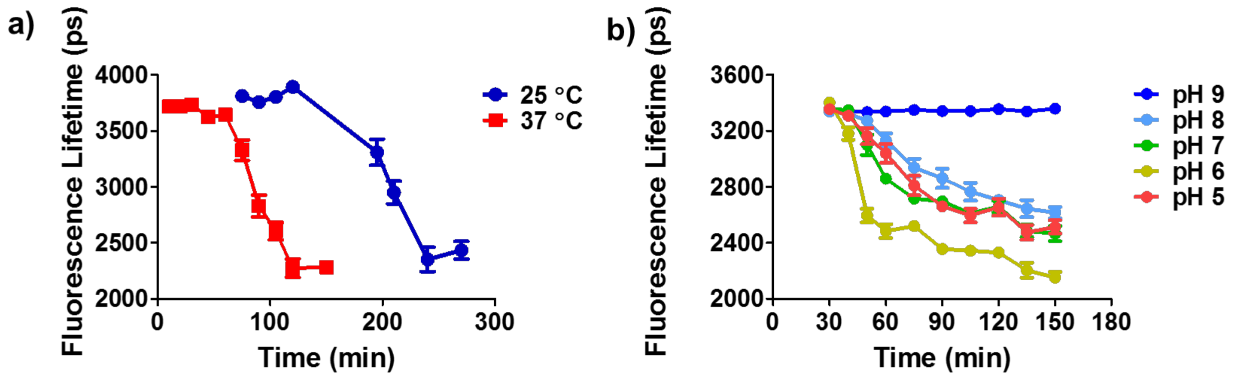

**Figure S2: Monitoring the effect of temperature and pH on $A\beta_{42}$ aggregation with the nanoFLIM**. **a)** Effect of temperature of aggregation profile**.** 10 µM $A\beta_{42}$, 50% labelled, n = 18.**b)** Effect of varying pH on aggregation profile. No aggregation was observed at pH 9 and an accelerated aggregation rate was observed at pH 6. 10 µM $A\beta_{42}$, 50% labelled, n = 8, TRIS buffer. Plots show mean ± SEM.

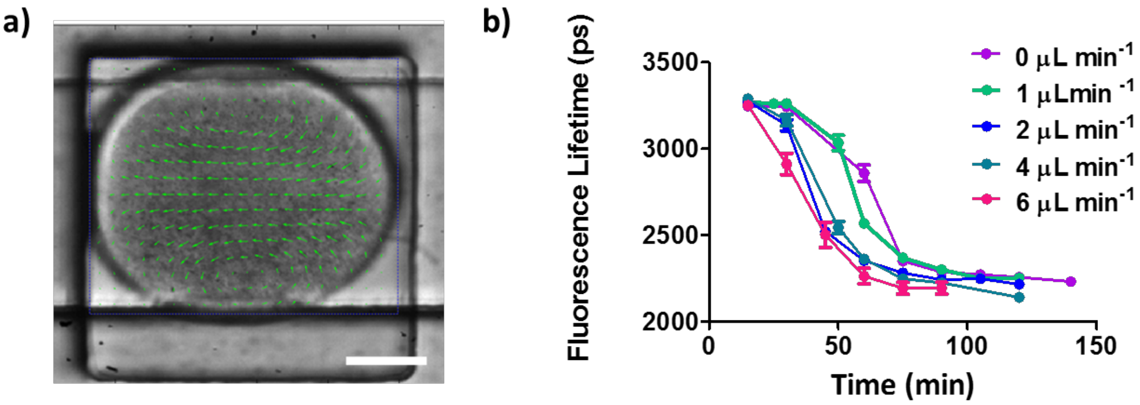

**Figure S3: The effect of shearing on the aggregation process.** a) A flow of oil under a trapped droplet can be used to generate a shear force, causing convection flows within the droplet. Green arrows obtained using Particle Image Velocimetry (PIV) methods indicate the amplitude and direction of flow of the droplet contents (see Supplementary Video 3). b) Increasing the oil flow rate under the trapped droplets results in an increased rate of aggregation, 1.75-fold rate increase at a 6 μL min-1 oil flow compared to static droplets. Plots show mean ± SEM, n= 20-30, 20 μM Aβ42, 50% labelled.

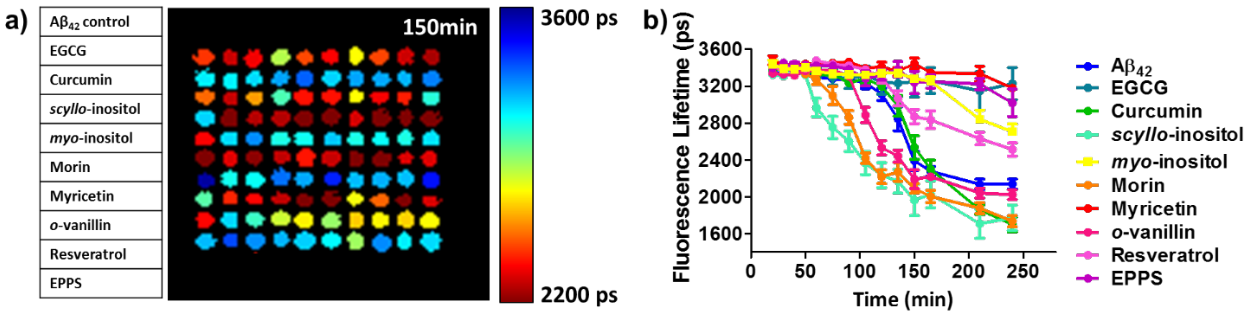

c)

| Compound | TEM | ThT Fluorescence | nanoFLIM | A11 |
|---|---|---|---|---|
| **EGCG**[1,2] | 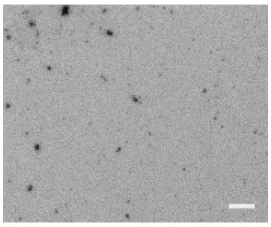 Small clumped structures | Strong Inhibition | Strong Inhibition | ✗ |
| **Curcumin**[3] | 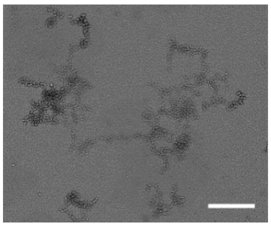 High density of curly fibrils | Inhibition | No inhibition - Reduced final fluorescence lifetime | ✓ |

| | | | | |
|---|---|---|---|---|
| ***scyllo*-inositol**[4] | 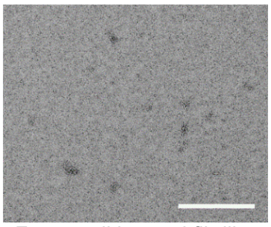 Few small knotted fibrillar species | No inhibition | No inhibition<br>- Accelerated aggregation | ✓ |
| ***myo*-inositol**[4] | 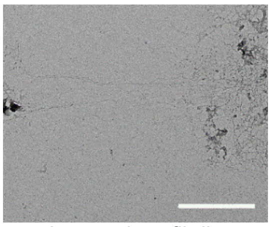 Long stringy fibrils | No inhibition<br>- No lag phase | Moderate inhibition<br>- Lengthened lag phase | ✗ |
| **Morin**[5] | 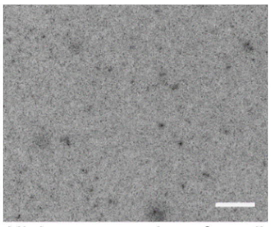 High concentration of small aggregates | Moderate inhibition | No inhibition<br>- Accelerated aggregation<br>- Reduced final fluorescence lifetime | ✓ |
| **Myricetin**[6] | 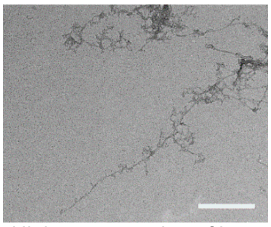 High concentration of long thin fibrils | Strong Inhibition | Strong Inhibition | ✓ |
| **o-Vanillin**[7] | 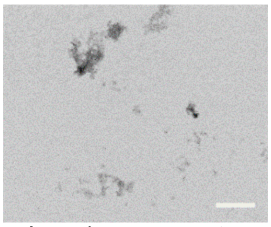 Amorphous aggregates | Moderate inhibition<br>- Reduced plateau | No inhibition<br>- Accelerated aggregation | ✓ |
| **Resveratrol**[8] | 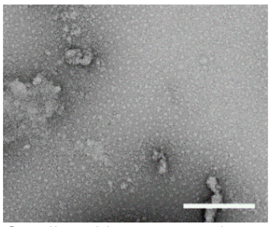 Small and large amorphous aggregates | Moderate inhibition<br>- no lag phase | Moderate inhibition | ✗ |

| | | | | |
|---|---|---|---|---|
| **EPPS**[9] | 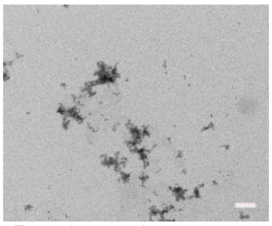 Few clustered aggregates | No inhibition | Strong Inhibition | ✗ |

**Figure S4: Detailed analysis of a collection of known inhibitors in the NanoFLIM assay and comparison to other widely used assays.**

a) The nanoFLIM was tested using a series of small molecules known to modulate the process of Aβ42 aggregation: EGCG, curcumin, scyllo-inositol, myo-inositol, morin, myricetin, o-vanillin, resveratrol and EPPS. Each row contains 10 droplets of 10 μM Aβ42 (50% labelled) with 50 μM of each of the small molecule modifiers, following 150 min incubation. b) Aβ42 aggregation kinetics obtained using the nanoFLIM. The blue curve shows the aggregation profile of Aβ42 in the absence of any small molecules. Plot shows mean ± SEM, n = 10. c) Comparison of the inhibitory activity of the small molecule observed with TEM, ThT, nanoFLIM and immunoassay (oligomer-specific A11 antibody) analysis. The nanoFLIM shows a higher success rate in correctly assigning the modulatory effect of small molecules on Aβ42 aggregation, as confirmed by TEM (Scale = 50 nm). Ticks (✓) are used to denote a positive A11 antibody result from Aβ42 samples incubated with the test compound, indicative of the presence of toxic oligomeric species. Crosses (✗) denote no A11 sensitivity.

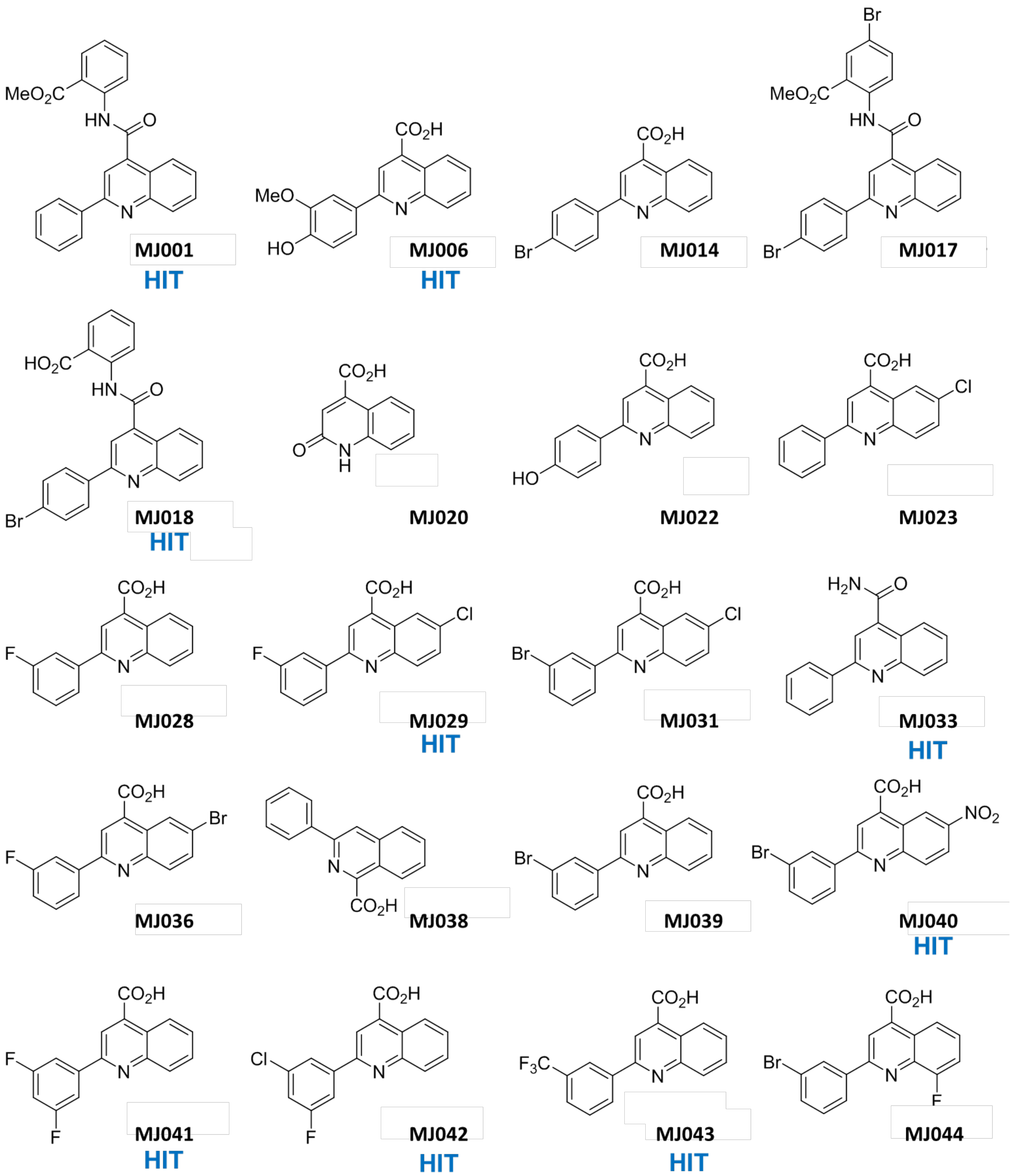

**Figure S5: Cinchophen derivative library.** Structures of the full cinchophen library. Compounds identified as inhibitors by nanoFLIM screening are marked with 'HIT'.[4]

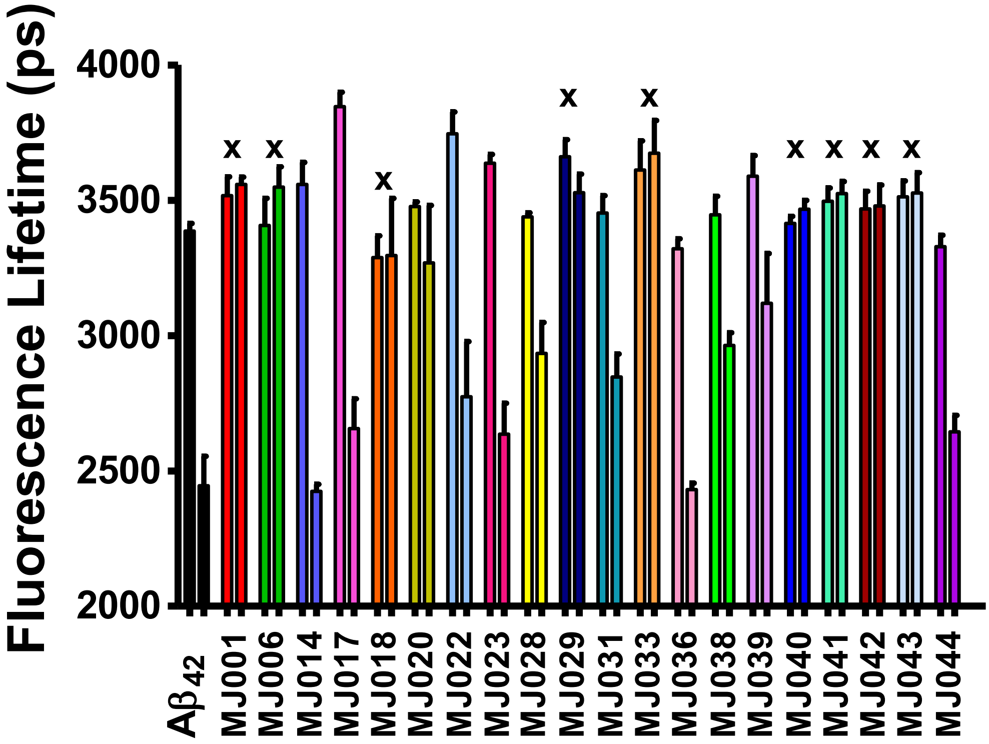

**Figure S6: Start and end points fluorescence lifetime values of Aβ42-488 incubated with each of the cinchophen library**. Fluorescence lifetime was monitored with nanoFLIM screening and values are given for each compound at t = 20 and t = 150 minutes. Compounds showing >30% inhibitory activity after 2.5 h incubation are marked with an 'X'. Plot shows mean + SEM for one of three independent repeats, n = 5 droplets, 10 µM $A\beta_{42}$,488, 50% labelled, 50 µM compound.

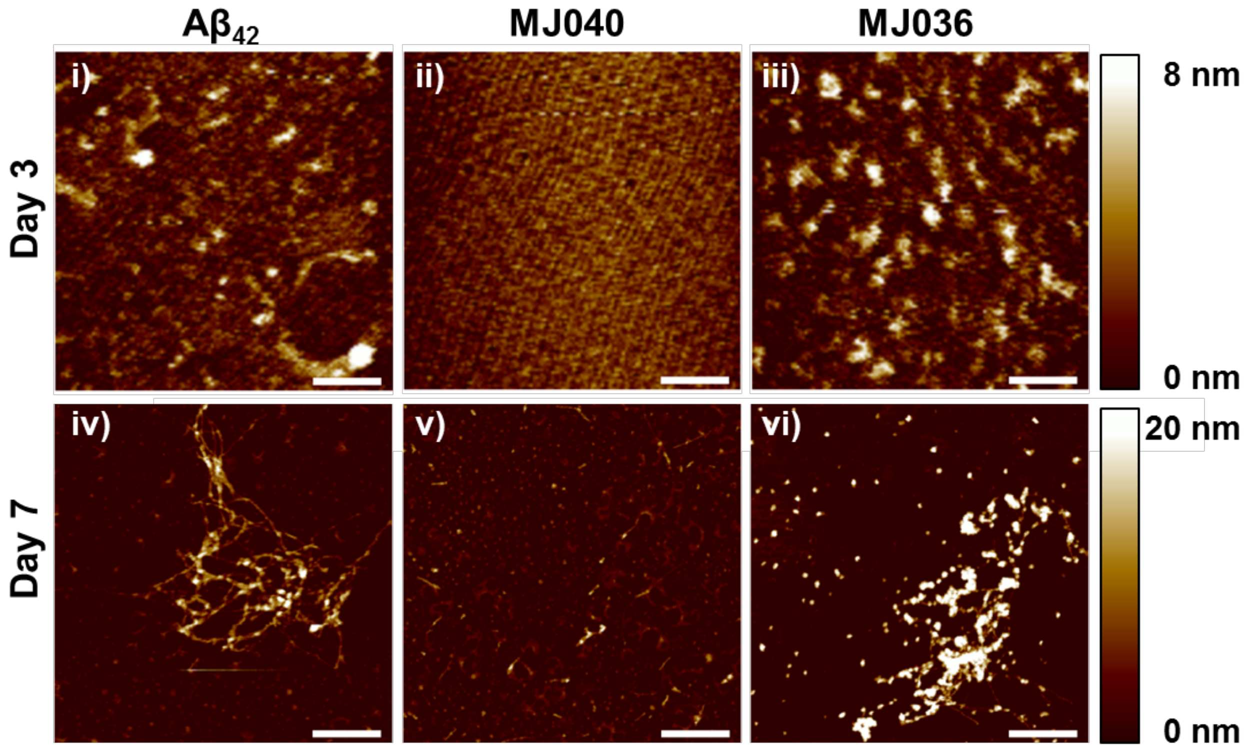

**Figure S7: AFM images of Aβ42 species formed in the absence or presence of a fivefold excess of compounds MJ036 or MJ040 following 3- or 7-day incubations**. A$\beta_{42}$ control; small aggregates observed at day 3 ***(i)*** and long amyloid fibrils present at day 7 ***(iv)***. **MJ040** treated; no structures observed at day 3 ***(ii)***, but some small species evident at day 7 ***(v)***. **MJ036** treated; at day 3 many aggregates are observed ***(iii)***. Longer fibrillar species with clumps along the structure are detected at day 7 ***(vi)***. Images were acquired with tapping mode AFM in air. ***i,ii,iii*** scale bar = 100 nm. i***v,v,vi*** scale bar = 600 nm.

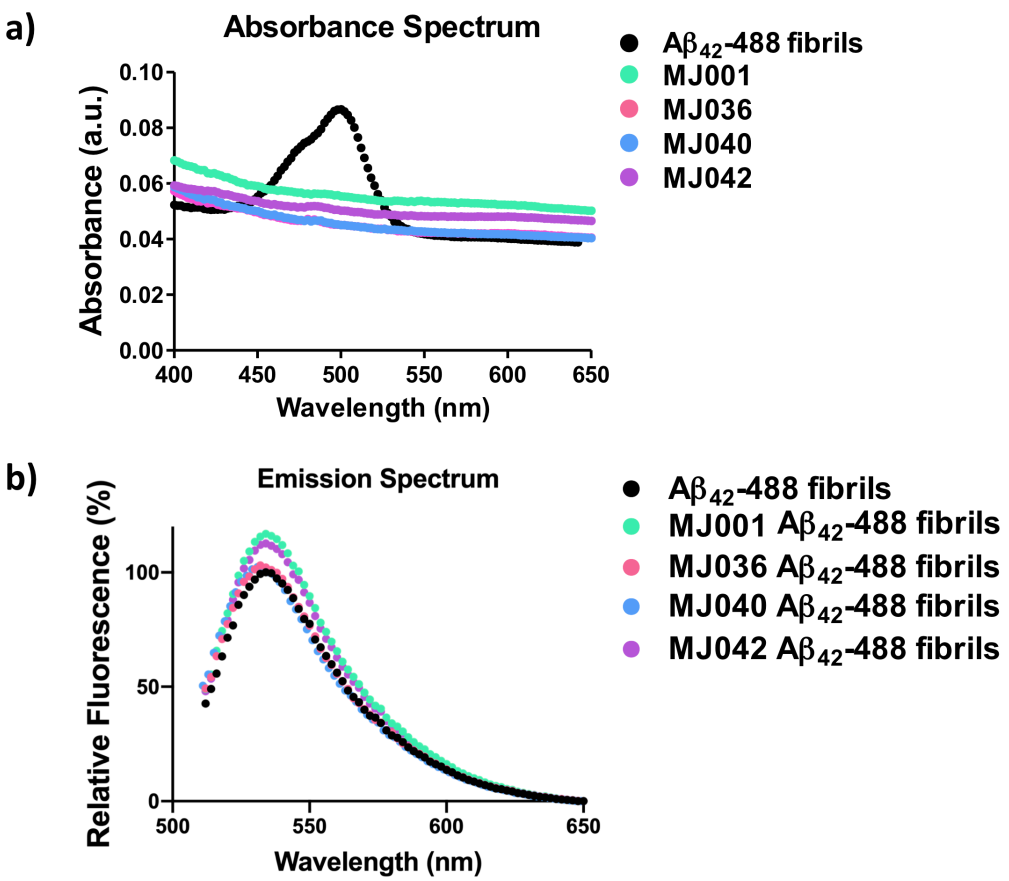

**Figure S8: Monitoring the fluorescence interference of select compounds at the excitation wavelength used in the nanoFLIM assay. a)** Comparison of the absorbance spectrum of A$\beta_{42}$-488 fibrils (10 µM, 50% labelled) and of 50 µM of each of the compound. The A$\beta_{42}$-488 fibrils give an absorbance spectrum characteristic for the dye. All compounds show lower absorbance. **b)** Emission spectrum of preformed A$\beta_{42}$-488 fibrils (10 µM, 50% labelled) in the presence of select compounds. The samples were excited at 480 nm, the wavelength used with the nanoFLIM assay. The fluorescence intensity of the partially labelled fibrils is unaffected by the presence of **MJ001, MJ036**, **MJ040** and **MJ042.**

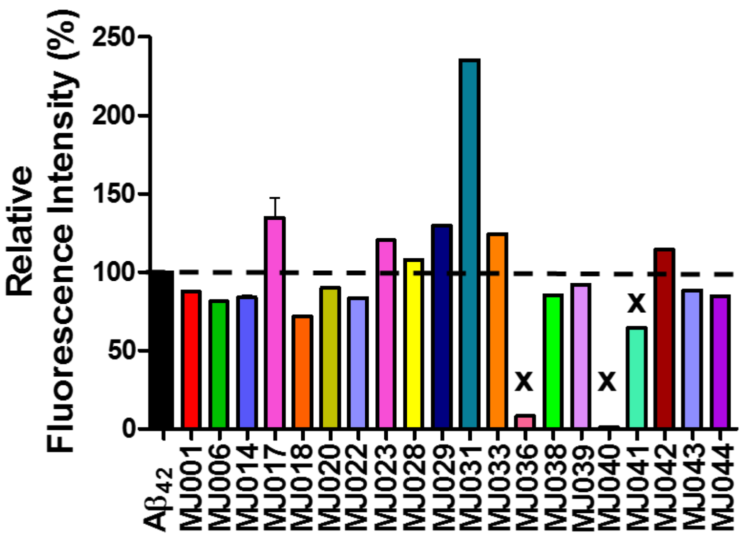

**Figure S9: Relative changes in the Aβ42-ThT fluorescence intensity at the aggregation plateau following 6 h of incubation with each of the compounds of the cinchophen library**. The ThT fluorescence in the untreated A$\beta_{42}$ sample is set as 100% and taken to represent fibril mass concentration of untreated A$\beta_{42}$. Compounds showing >30% inhibitory activity after 6 h incubation are marked with an 'X' (**MJ036** and **MJ040**). Plot shows mean + SEM, n = 3, 10 µM A$\beta_{42}$, 20 µM ThT, 50 µM compound.

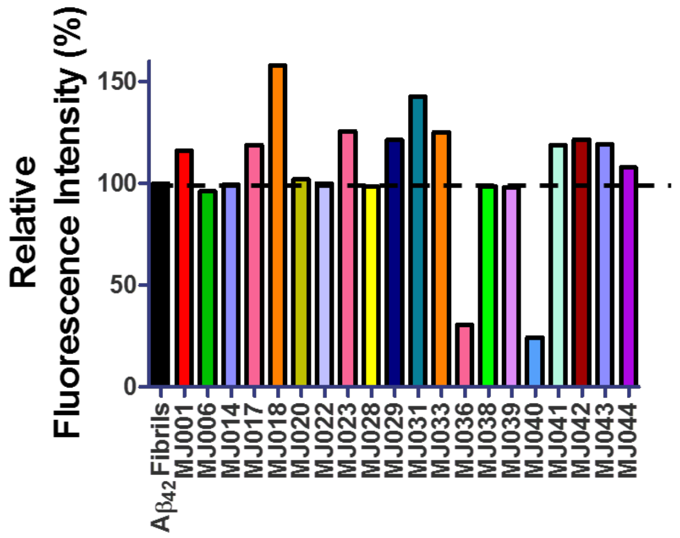

**Figure S10: Relative changes in the intensity of the ThT fluorescence emission spectra of preformed Aβ42 fibrils with addition of cinchophen library compounds.** The fluorescence intensity of the ThT-$A\beta_{42}$ fibril sample is set to 100%. Addition of compounds **MJ001**, **MJ017**, **MJ018**, **MJ023**, **MJ029**, **MJ031, MJ041, MJ042** and **MJ043** results in an increase in fluorescence intensity relative to the fibrils and ThT alone. The addition of **MJ036** and **MJ040** results in lowered fluorescence intensity, indicating interference by means of inner filter effects or competitive binding to ThT or ThT binding sites.[5] Intensity measured at 488 nm emission, with 440 nm excitation. 10 µM preformed $A\beta_{42}$ fibrils, 20 µM ThT, 50 µM compound.

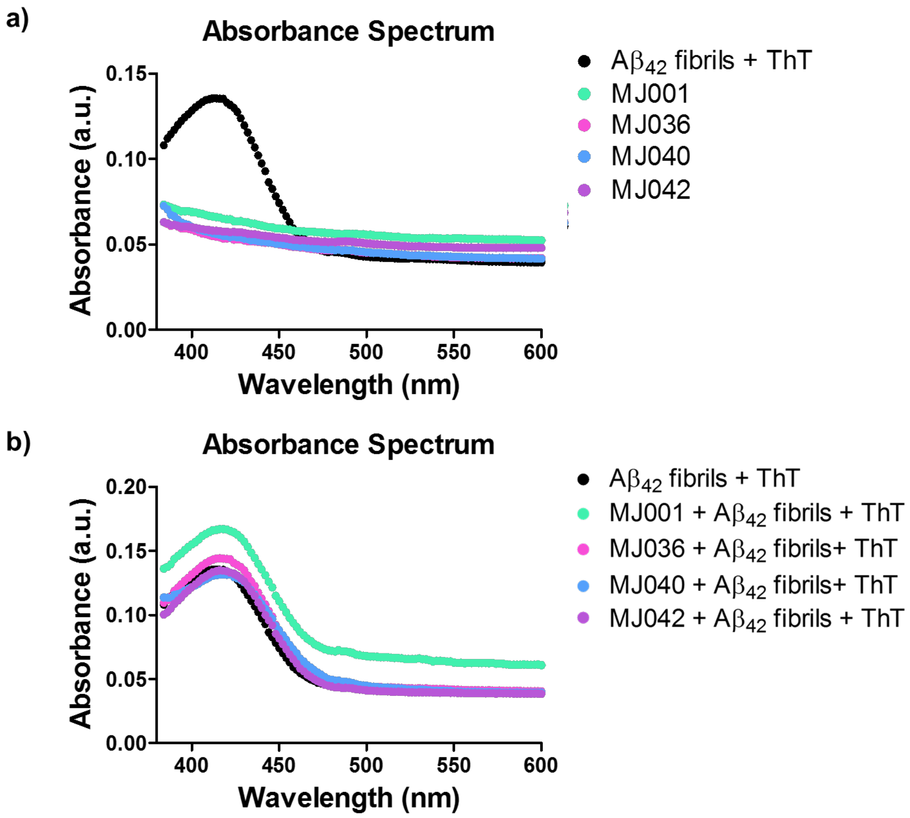

**Figure S11: Absorption spectra of the compounds compared to that of preformed Aβ42 fibrils with ThT.** a) The ThT-$A\beta_{42}$ fibril absorbance at 440 nm, the excitation wavelength used in the ThT fluorescence assay, dominates greatly over each of the individual compounds. 10 µM $A\beta_{42}$ fibrils, 20 µM ThT, 50 µM compound. **b)** The ThT-$A\beta_{42}$ fibril absorbance at 440 nm is largely unaffected by the addition of **MJ036**, **MJ040** and **MJ042.** The addition of MJ001 significantly increases the observed ThT absorbance.

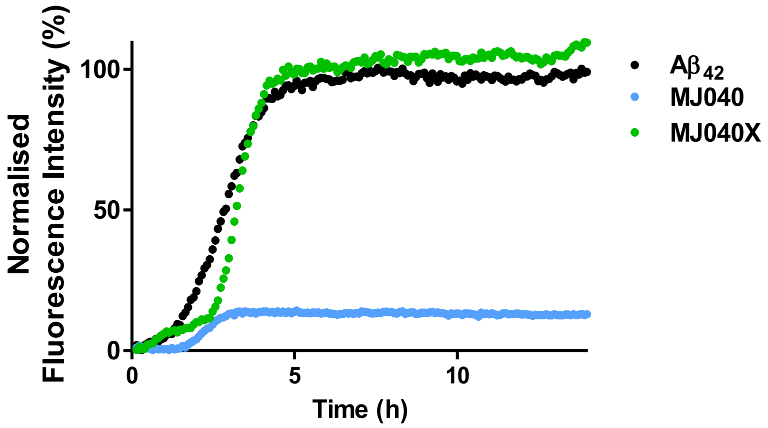

**Figure S12: Comparison of the *in vitro* inhibitory activity of MJ040 and MJ040X by means of ThT fluorescence.** The curve for Aβ42 (black) represents the time course of Aβ42 aggregation in the absence of inhibitors, the plateau of which is taken to represent fibril mass concentration and is set as 100%. **MJ040X** shows little effect on the *in vitro* aggregation of Aβ42, in contrast to **MJ040**, which shows a large inhibitory effect**.** Aβ42 10 µM Aβ42, 50 µM compound, 20 µM ThT.

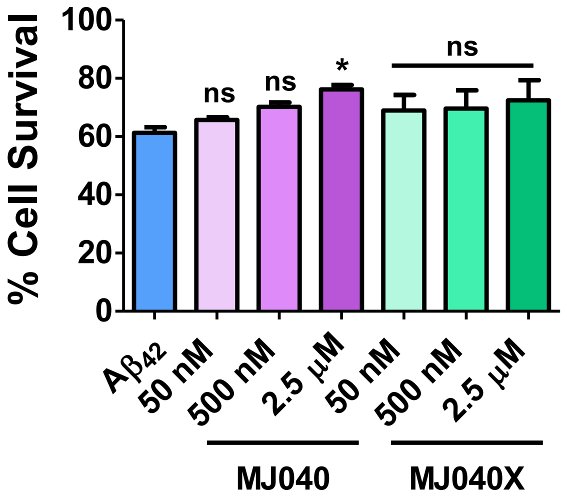

**Figure S13: Monitoring the rescuing effect of MJ040 and MJ040X from pre-aggregated Aβ42 induced toxicity.** Aβ42 (10 µM) was pre-incubated for 24 h with or without the compounds, then diluted (500 nM) and added to SH-SY5Y cells. After 48 h treatment, cytotoxicity was evaluated using an MTT assay. The species formed in the presence in the active drug **MJ040** induced less cell death than the untreated peptide. **MJ040X** has poor *in vitro* inhibitory activity and did not rescue the cells from pre-aggregated Aβ42 induced cytotoxicity. The viability of untreated cells was set as 100%. Error bars represent SEM, n = 4, statistical analysis performed using one-way ANOVA with Dunnett's multiple comparison post-test; *$p<0.05$.

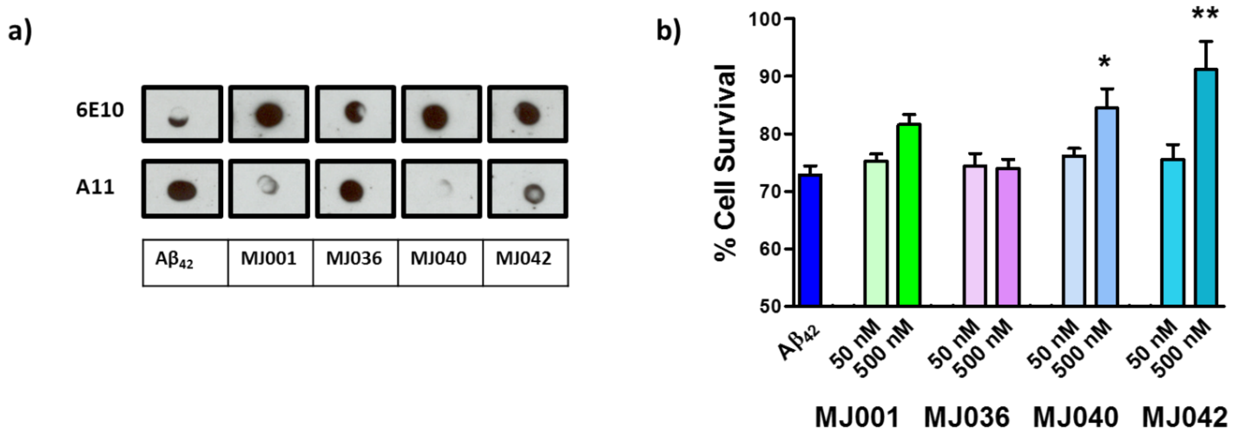

**Figure S14: Validation of inhibitory activity of subset of cinchophen compounds. a)** Dot blots to investigate the potential formation of oligomeric $A\beta_{42}$ using oligomer-specific antibody A11, following 5-day incubation with **MJ001**, **MJ036, MJ040** or **MJ041**. A11-sensitive species were detected in the $A\beta_{42}$ control or when the peptide was incubated in the presence of **MJ036**. Positive readouts were obtained for all samples using the 6E10 control antibody, which detects all Aβ species. **b)** Monitoring the rescuing effect of **MJ001**, **MJ036**, **MJ040** and **MJ042** from monomeric $A\beta_{42}$ induced toxicity. SH-SH5Y cells were incubated in the compound for 6 h prior to the addition of $A\beta_{42}$ monomers (250 nM). Following 48 h incubation the percentage of cell survival was assessed using an MTT cell viability assay. **MJ040** (500 nM) and **MJ042** (500 nM) were seen to significantly increase the cell viability following exposure to $A\beta_{42}$. The viability of untreated cells was set as 100%. Error bars represent SEM, n = 4, statistical analysis performed by one-way ANOVA with Dunnett's multiple comparison post-test; $^*p<0.05$, $^{**}p<0.01$.

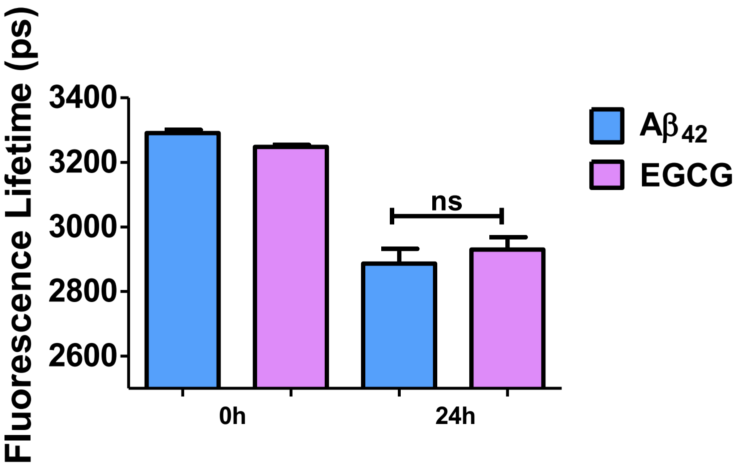

**Figure S15: Cellular fluorescence lifetime screen with EGCG addition at the same time as $A\beta_{42}$-488.** If the labelled peptide and inhibitory EGCG are added to the extracellular medium at the same time, there is no significant effect on intracellular $A\beta_{42}$ aggregation. 250 nM $A\beta_{42}$, 50% labelled, 24 h incubation, mean lifetime + SEM, n = 8-10, statistical analysis performed using a Student's t-tests.

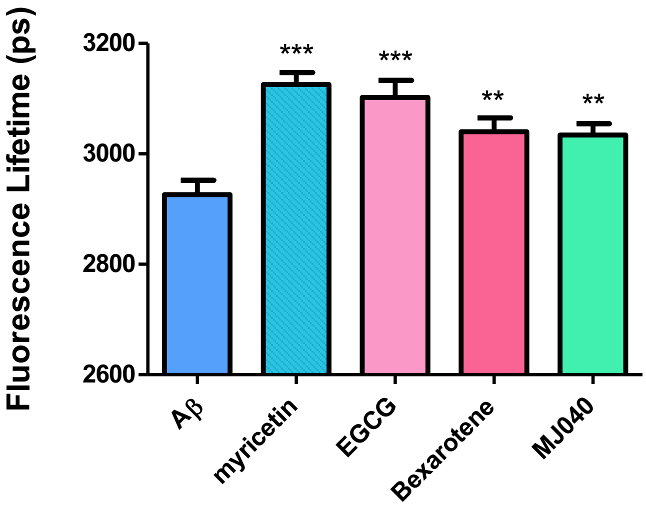

**Figure S16: Validation of the cellular fluorescence lifetime assay with known inhibitors.** The cellular fluorescence lifetime assay was validated with known small molecules inhibitors myricetin,[6] EGCG[7, 8] and bexarotene.[9] These were all shown to inhibit the aggregation of internalised labelled Aβ42, relative to the untreated control. Original hit compound **MJ040** also showed anti-aggregation activity. 250 nM Aβ42, 50% labelled, 24 h incubation, mean lifetime + SEM, n = 25-30. Statistical significant analysed using a one-way ANOVA with Dunnett's multiple comparison post-test; * = $p<0.05$, ** = $p<0.01$, ***=$p<0.001$.

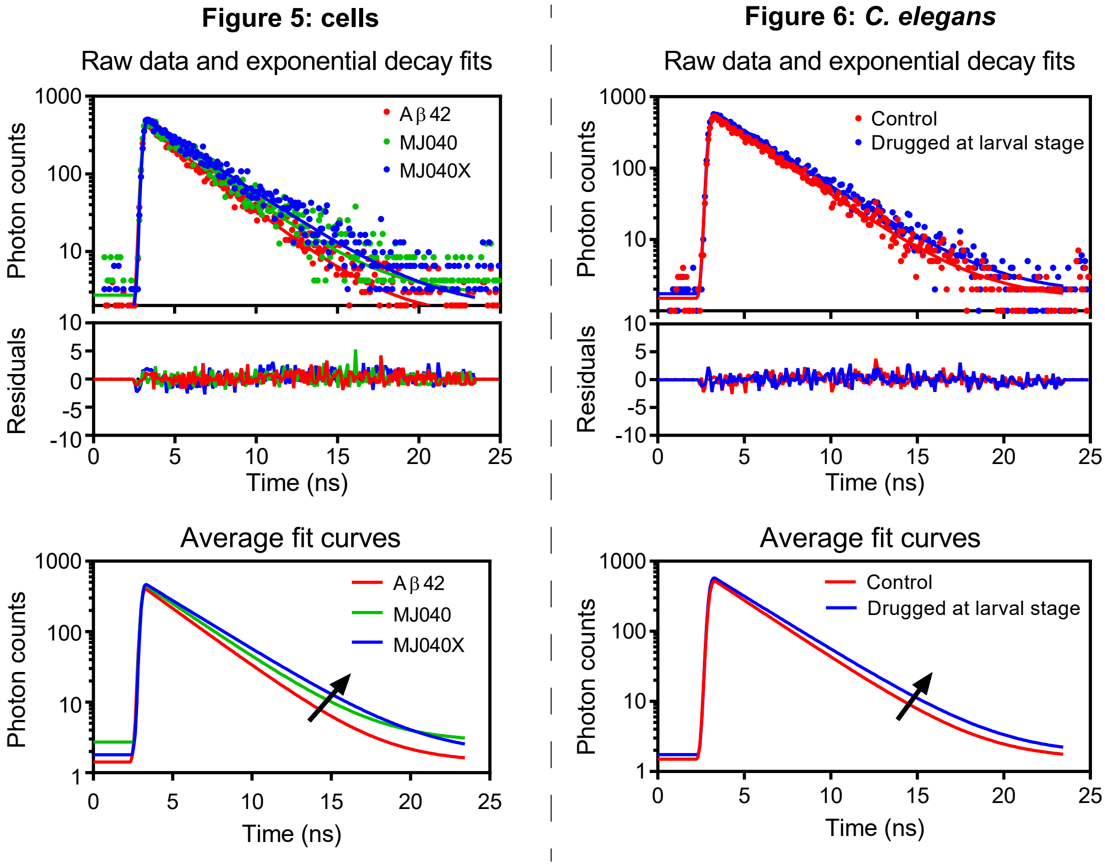

**Figure S17: Time-correlated single-photon counting (TCSPC).** Typical TCSPC time traces and exponential decay fits for Aβ42-488 cell data in Figure 5 and GFP-Aβ42 *C. elegans* data in Figure 6. Time traces were fitted to a single exponential decay.

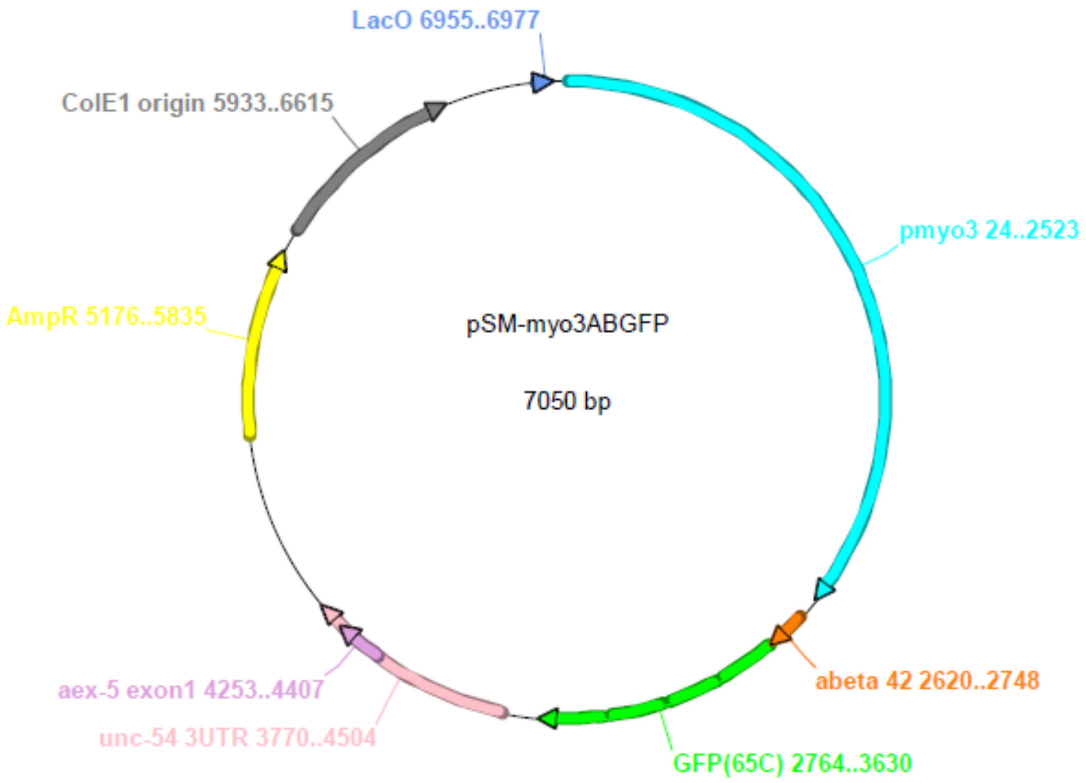

**Figure S18: Plasmid map of the construct Pmyo3::GFP::Aβ42 used for GFP expression in *C. elegans*.** GFP and Aβ42 were under control of the promoter pmyo3 and expressed in muscle myosin. We thank Dr Kaveh Ashrafi and Jason Liu for making the *C. elegans* strain containing this plasmid available to us.

### Supplementary Tables

**Table S1: Compound libraries tested in this pilot nanoFLIM screening campaign**

| Library Name | Number screened | Number of hits* | Study Title | Reference |
|---|---|---|---|---|
| Partially Saturated Bicyclic Heteroaromatics Fragments | 21 | 5 | Partially Saturated Bicyclic Heteroaromatics as an $sp^3$ -Enriched Fragment Collection | 10 |
| Quinolone Natural Products derivatives | 7 | 0 | Divergent Synthesis of Quinolone Natural Products from Pseudonocardia sp. CL38489 | 11 |
| Unpublished DOS library | 10 | 0 | Unpublished | — |
| DOS drug like macrocycles | 25 | 3 | Diversity-Oriented Synthesis of Drug-Like Macrocyclic Scaffolds Using an Orthogonal Organo- and Metal Catalysis Strategy | 12 |
| Quinolizin-4-ones and pyrido[1,2-a]pyrimidin-2-ones | 44 | 3 | Concise Synthesis of Substituted Quinolizin-4-ones by Ring-Closing Metathesis<br><br>Concise synthesis of rare pyrido[1,2-a]pyrimidin- 2-ones and related nitrogen-rich bicyclic scaffolds with a ring-junction nitrogen[17] | 13<br><br>14 |
| Paracyclophane analogues | 28 | 0 | Unpublished | |
| Flavone + heterocyclic flavone derivatives | 54 | 2 | Divergent and concise total syntheses of dihydrochalcones and 5-deoxyflavones recently isolated from Tacca species and Mimosa diplotricha<br><br>Divergent Total Syntheses of Flavonoid Natural Products Isolated from Rosa rugosa and Citrus unshiu | 15<br><br>16 |
| Flavone dimers | 30 | 0 | Divergent synthesis of biflavonoids yields novel inhibitors of the aggregation of amyloid β (1–42) | 17 |

| Library Name | Number screened | Number of hits* | Study Title | Reference |
|---|---|---|---|---|
| Chalcones and heteroaromatic chalcone derivatives | 32 | 19 | Unpublished | — |
| Triazole linked flavonoid derivatives | 31 | 1 | Combinatorial Synthesis of Structurally Diverse Triazole-Bridged Flavonoid Dimers and Trimers | 18 |
| Aurones + biaurones | 32 | 14 | Divergent synthesis of biflavonoids yields novel inhibitors of the aggregation of amyloid β (1–42)[20] | 17 |
| Build/Couple/Pair/Diversify Macrocylic DOS library | 111 | 10 | A strategy for the diversity-oriented synthesis of macrocyclic scaffolds using multidimensional coupling[22]<br><br>A Multidimensional Diversity-Oriented Synthesis Strategy for Structurally Diverse and Complex Macrocycles[23] | 19<br><br>20 |
| Phenyl Quinoline library | 20 | 9 | Allosteric modulation of AURKA kinase activity by a small-molecule inhibitor of its protein-protein interaction with TPX2[10] | 4 |

**Table S2: Statistical analysis for Figure 5a.** Statistical analysis of the difference in the mean fluorescence lifetime (ps) in control **A$\beta_{42}$** treated cells, and cells treated with **ECGC** (250nM and 2.5nM). P-values were computed in GraphPad Prism using a two-way Anova and a Bonferroni post-test. ***p<0.001.

a)

| Two-way ANOVA with Bonferroni post-test | | | | | | | |
|---|---|---|---|---|---|---|---|
| | **Control A$\beta_{42}$** | **250 nM ECGC** | ***Difference*** | ***95% CI of diff.*** | **t** | **P value** | **Summary** |
| 0 | 3299 | 3342 | 43.21 | -186.5 to 272.9 | 0.5841 | P > 0.05 | ns |
| 12 | 2759 | 2938 | 179.4 | 66.59 to 292.2 | 4.938 | P < 0.001 | *** |
| 24 | 2827 | 2992 | 165.2 | 38.31 to 292.0 | 4.043 | P < 0.001 | *** |
| 48 | 2834 | 2988 | 153.3 | 22.86 to 283.7 | 3.650 | P < 0.01 | ** |

b)

| **Time (hr)** | **Control A$\beta_{42}$** | **2.5 µM ECGC** | ***Difference*** | ***95% CI of diff.*** | **t** | **P value** | **Summary** |
|---|---|---|---|---|---|---|---|
| 0 | 3299 | 3329 | 30.25 | -215.3 to 275.8 | 0.3825 | P > 0.05 | ns |
| 12 | 2759 | 2996 | 237.2 | 116.2 to 358.3 | 6.088 | P<0.001 | *** |
| 24 | 2827 | 3055 | 228.3 | 106.6 to 350.1 | 5.823 | P<0.001 | *** |
| 48 | 2834 | 3025 | 190.3 | 72.31 to 308.2 | 5.009 | P<0.001 | *** |

c)

| **Time (hr)** | **250 nM ECGC** | **2.5 µM ECGC** | ***Difference*** | ***95% CI of diff.*** | **t** | **P value** | **Summary** |
|---|---|---|---|---|---|---|---|
| 0 | 3342 | 3329 | -12.96 | -242.7 to 216.7 | 0.1752 | P > 0.05 | ns |
| 12 | 2938 | 2996 | 57.83 | -65.92 to 181.6 | 1.451 | P > 0.05 | ns |
| 24 | 2992 | 3055 | 63.15 | -60.06 to 186.4 | 1.592 | P > 0.05 | ns |
| 48 | 2988 | 3025 | 36.99 | -93.42 to 167.4 | 0.8809 | P > 0.05 | ns |

**Table S3: Statistical analysis for Figure 5b.** Statistical analysis of the difference in the mean fluorescence lifetime (ps) in control **A$\beta_{42}$** treated cells, **MJ040** treated cells, and MJ040X treated cells. P-values were computed in GraphPad Prism using one-way ANOVA with Tukey's test for multiple comparisons. ****p<0.0001.

| One-way ANOVA with Tukey's Multiple Comparison post-test | | | | | | |
|---|---|---|---|---|---|---|
| | **Means** | **Mean Diff.** | **q** | **Significant? P < 0.05?** | **Summary** | **95% CI of diff** |
| A$\beta_{42}$-488 *vs* A$\beta_{42}$-488+MJ040 | 2368 *vs* 2616 | -247.8 | 10.26 | Yes | **** | -328.6 to -167.0 |
| A$\beta_{42}$-488 *vs* A$\beta_{42}$-488 +MJ040X | 2368 *vs* 2874 | -506.1 | 19.77 | Yes | **** | -591.7 to -420.5 |
| MJ040 *vs* MJ040X | 2616 *vs* 2874 | -258.3 | 9.755 | Yes | **** | -346.9 to -169.8 |

**Table S4: Statistical analysis for Figure 6d.** Statistical analysis of the difference in the mean fluorescence lifetime (ps) of untreated control GFP:Aβ worms and the **MJ040X** treated worms. P-values were computed in GraphPad Prism using the either a two-way Anova and a Bonferroni post-test or a one-way Anova and a Dunnett's test of multiple comparison. *p<0.05; **p<0.01, ***p<0.001.

**a)**

| Two-way ANOVA with Bonferroni post-test | | | | | | | |
|---|---|---|---|---|---|---|---|
| ***Day*** | ***Control*** | ***MJ040X*** | ***Difference*** | ***95% CI of diff.*** | t | P value | Summary |
| Day 1 | 2769 | 2769 | 0 | -43.45 to 43.45 | 0 | P > 0.05 | ns |
| Day 3 | 2754 | 2747 | -6.824 | -51.53 to 37.89 | 0.4407 | P > 0.05 | ns |
| Day 6 | 2745 | 2743 | -2.163 | -47.56 to 43.24 | 0.1375 | P > 0.05 | ns |
| Day 9 | 2724 | 2745 | 21.49 | -22.74 to 65.71 | 1.403 | P > 0.05 | ns |
| Day 12 | 2698 | 2754 | 56.56 | 5.635 to 107.5 | 3.207 | P<0.01 | ** |
| Day 15 | 2715 | 2714 | -1.646 | -42.37 to 39.08 | 0.1167 | P > 0.05 | ns |

**b)**

| One-way ANOVA with Dunnett's Multiple Comparison Test | | | | | |
|---|---|---|---|---|---|
| | **Mean Diff.** | **q** | **Significant? P < 0.05?** | **Summary** | **95% CI of diff** |
| Control Day 1 *vs* Control Day 3 | 15.01 | 1.086 | No | ns | -23.41 to 53.44 |
| Control Day 1 *vs* Control Day 6 | 23.99 | 1.708 | No | ns | -15.05 to 63.03 |
| Control Day 1 vs Control Day 9 | 45.33 | 3.371 | Yes | ** | 7.958 to 82.70 |
| Control Day 1 *vs* Control Day 12 | 71.44 | 4.431 | Yes | *** | 26.62 to 116.2 |
| Control Day 1 *vs* Control Day 15 | 53.6 | 4.036 | Yes | *** | 16.68 to 90.51 |
| Control Day 1 *vs* MJ040X Day 1 | 0 | 0 | No | ns | -37.87 to 37.87 |
| Control Day 1 *vs* MJ040X Day 3 | 21.84 | 1.58 | No | ns | -16.59 to 60.26 |
| Control Day 1 *vs* MJ040X Day 6 | 26.16 | 1.892 | No | ns | -12.27 to 64.58 |
| Control Day 1 *vs* MJ040X Day 9 | 23.84 | 1.698 | No | ns | -15.20 to 62.88 |
| Control Day 1 *vs* MJ040X Day 12 | 14.88 | 1.106 | No | ns | -22.49 to 52.25 |
| Control Day 1 *vs* MJ040X Day 15 | 55.24 | 4.207 | Yes | *** | 18.75 to 91.74 |

**Table S5: Statistical analysis for Figure 6e** Statistical analysis of the difference in the mean fluorescence lifetime (ps) of the untreated control GFP:Aβ worms and the worms treated with **MJ040X** from larvae. P-values were computed in GraphPad Prism using a one-way Anova and a Sidak's test for multiple comparisons. **p<0.01.

| One-way ANOVA with Sidak's Multiple Comparison Test | | | | | | |
|---|---|---|---|---|---|---|
| | **Means** | **Mean Diff.** | **q** | **Significant? P < 0.05?** | **Summary** | **95% CI of diff** |
| Control Day 1 *vs* Adult MJ040X Day 1 | 2764 *vs* 2770 | -5.96 | 0.3804 | No | ns | -43.68 to 31.76 |
| Control Day 6 *vs* Adult MJ040X Day 6 | 2747 *vs* 2768 | -20.86 | 1.425 | No | ns | -56.12 to 14.39 |
| Control Day 15 *vs* Adult MJ040X Day 15 | 2722 *vs* 2780 | -58.04 | 3.538 | Yes | ** | -97.53 to -18.55 |

### Synthetic procedures

**Synthesis of MJ040 - (2-(3-bromophenyl)-6-nitroquinoline-4-carboxylic acid**

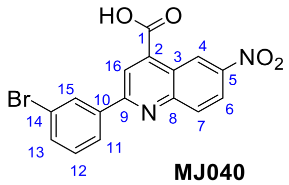

**MJ040**

According to the procedure described by Giardina *et al.*,[21] 5-nitroisatin (1.00 g, 5.21 mmol), 3'-bromoacetophenone (1.25 g, 6.25 mmol) and KOH (876 mg, 15.62 mmol) were dissolved in EtOH (80 mL). The reaction mixture was heated under reflux for 48 h, allowed to cool, then the solvent removed under reduced pressure. The residue was dissolved in $H_2O$ (100 mL) and washed with $Et_2O$ (100 mL x 3). The aqueous layer was cooled to 0 °C, acidified to pH 2 with 3 M HCl, and the precipitate was filtered under suction. The crude residue was purified by column chromatography over silica (10% MeOH in $CH_2Cl_2$) to afford a light brown solid (110 mg, 0.42 mmol, 8%). **TLC** $R_f$ = 0.12 (10% MeOH in $CH_2Cl_2$); **IR** $\nu_{max}$ (neat)/cm$^{-1}$: 2923 m (C-H), 1706 m (C=O), 1588 s (C=C), 1528 m (N=O), 1328 m (N=O), 1223 m, 1056 ; **$^{1}$H NMR** (500 MHz, DMSO): δ 7.56 (1H, d, *J* = 8.0 Hz, ArH12), 7.79 (1H, ddd, *J* = 8.0, 2.0, 0.9 Hz, ArH11/ArH13), 8.36 (1H, ddd, *J* = 8.0, 1.5, 0.9 Hz, ArH11/ ArH13), 8.38 (1H, d, *J* = 9.3 Hz, ArH7), 8.52-8.57 (2H, m, ArH6, ArH15), 8.70 (1H, s, ArH16), 9.66 (1H, d, *J* = 2.5, ArH4) ppm; **$^{13}$C NMR** (126 MHz, DMSO): δ 121.6 (ArC16), 122.7 (ArC4), 123.0 (Cq), 123.7 (ArC6), 126.9 (ArC11/ArC13), 130.2 (ArC15), 131.4 (C12), 131.8 (ArC7), 133.7(ArC11/ArC13), 139.1 (Cq), 139.4 (Cq), 146 (Cq), 150.4 (Cq), 152.8 (Cq) 157.9 (Cq), 166.7 (C=O) ppm; **HRMS** (ESI) *m/z* = 372.9810, $[M+H]^+$ found, $C_{16}H_{10}O_4N_2{}^{79}B^+$ required 372.9818.

**Synthesis of MJ040X - methyl 2-(3-bromophenyl)-6-nitroquinoline-4-carboxylate**

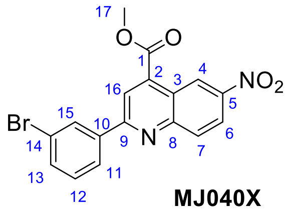

**MJ040X**

To a mixture of 2-(3-bromophenyl)-6-nitroquinoline-4-carboxylic acid **MJ040** (100 mg, 0.268 mmol) in MeOH (10 mL) was added conc. $H_2SO_4$. The mixture was heated under reflux for 24 h, then allowed to cool and neutralized with NaOH (25%). The solvent was reduced under vacuum. EtOAc (20 mL) was added and the organic layer was washed with saturated $NaHCO_3$ (2 × 20 mL) and brine (2 × 20 mL) then dried over anhydrous $MgSO_4$. The crude residue was purified by flash column chromatography over silica with a 0-50% gradient EtOAc in Hexane to afford a pale orange solid **MJ040X** (71.1 mg, 0.182 mmol, 68% yield). **TLC** $R_f$ = 0.83 (1:1 EtOAc/Hex); **m.p.** 164-165 °C; **IR** $\nu_{max}$ (neat)/cm$^{-1}$: 2962 m (C-H), 1724 s (C=O), 1530 s (N=O), 1501 m, 1436 s, 1342 (N=O), 1269 s, 1077 s; **$^1$H NMR** (500 MHz, DMSO): δ 4.06 (3H, s, -$OCH_3$), 7.56 (1H, d, *J* = 7.9 Hz, ArH12), 7.79 (1H, ddd, *J* = 7.9, 1.9, 1.0 Hz, ArH11/ ArH13), 8.34 (1H, ddd, *J* = 1.0, 1.9, 7.9 Hz, ArH11/ ArH13), 8.36 (1H, dd, *J* = 9.2, 0.3 Hz, ArH7), 8.52 (1H, t, *J* = 1.8 Hz, ArH15), 8.54 (1H, d, *J* = 2.6, 9.2 Hz, ArH6), 8.69 (1H, s, ArH16), 9.53 (1H, d, *J* = 2.6, ArH4) ppm; **$^{13}$C NMR** (126 MHz, DMSO): δ 53.4 (-$OCH_3$), 121.6 (C16), 122.3 (C4), 122.6 (Cq), 122.7 (Cq), 123.9 (Cq), 126.9 (C11/C13), 130.2 (C15), 131.4 (C12), 131.8 (C7), 133.8 (C11/C13), 137.7(Cq) , 139.2 (Cq), 146.1 (Cq), 150.2 (Cq), 157.2 (Cq), 165.4 (C=O) ppm; **HRMS** (ESI) *m/z* = 386.9962 $[M+H]^+$ found, $C_{17}H_{12}N_2O_4Br^+$ required 386.9980.

**Supplementary calculations of materials used in different assay formats and their estimated costs**

**(A) General assumptions and study design**

- Total compounds = 445
- Five repeats per compound, one A$\beta_{42}$ control (5 repeats) per chip/plate
- Average molecular weight per compound = 400 g/mol
- Molecular weight: A$\beta_{42}$ 4514.1 g/mol, A$\beta_{42}$-488 4870.5 g/mol
- Average mass 50% labelled A$\beta_{42}$:
  [4514.1 g/mol (A$\beta_{42}$) + 4870.5 g/mol A$\beta_{42}$-488]/2 = 4692.3 g/mol
- Cost: A$\beta_{42}$ at the price of £215 per mg, A$\beta_{42}$-488 at the price of £183 per 100 µg (Eurogentec, UK)

**(B) Estimation of amounts needed for a NanoFLIM campaign**

Calculations were based on the following assumptions:

- Average compounds tested per screen = 20
- Average time to reach end point = 3 h
- Total assay time= 445/20 = 22.25 assays x 3 h = ~ 70 h
- Total experiments = 110 droplets per chip x 22.25 assays = ~ 2450 experiments

***Mass of peptide used***

i) Per droplet – 18 nL at 10 µM → 1.8 x$10^{-13}$ mol → 845 pg
ii) Per sample (including dead volume) – 10 µL at 10 µM → 1.1x$10^{-10}$ mol → 469 ng
iii) Entire screen – 22.25 assays x 21 samples* x 469 ng = 219 µg

***Cost of peptide used***

i) Per droplet - 845 pg A$\beta_{42}$-488 (50% labelled)
422.5 pg A$\beta_{42}$-488 at the price of £183 per 100 µg
422.5 pg A$\beta_{42}$ at the price of £215 per mg
→ 0.087 p

ii) Per sample - 469 ng A$\beta_{42}$-488 (50% labelled)
234.5 ng A$\beta_{42}$-488 at the price of £183 per 100 µg
234.5 ng A$\beta_{42}$ at the price of £215 per mg
→ 42.9 p + 5 p = £0.48

iii) Entire screen – 22.25 assays x 21 samples* x £0.48
→ £226.80

(*21 samples = 20 compounds + one A$\beta_{42}$ control)

***Mass of compound used***

i) Per droplet – 18 nL at 50 µM → 9 x$10^{-13}$ mol → 360 pg

ii) Per sample (including dead volume) – 10 µL at 50 µM → 5x$10^{-10}$ mol → 500 ng

**(C) Estimation of amounts needed for a ThT Fluorescence Assay campaign**

Calculations were based on the following assumptions:

- Average compounds tested per screen = 18 (5 replicates)
- Average time to reach end point = 8 h

Total assay time= 445/18 = 24.7 assays x 8 h = ~ 200 h

***Mass of peptide used***

i) Per well – 100 µL at 10 µM → 1x$10^{-9}$ mol → 4.5 µg
ii) Entire screen – 24.7 x 96-well plates x 4.5 µg = 10.7 mg

***Cost of peptide used***

i) Per well – 4.5 µg $A\beta_{42}$ at the price of £215 per mg
→ £0.97
ii) Entire screen – 10.7 mg $A\beta_{42}$ at the price of £215 per mg
→ £2300.5

***Mass of compound used***

i) Per well – 100 µL at 50 µM → 5x$10^{-9}$ mol → 200 ng
